# CRISPR gene and transcriptome engineering (CRISPRgate) improves loss-of-function genetic screening approaches

**DOI:** 10.1101/2024.05.14.594135

**Authors:** Jannis Stadager, Chiara Bernardini, Laura Hartmann, Henrik May, Jessica Wiepcke, Monika Kuban, Zeynab Najafova, Steven A. Johnsen, Stefan Legewie, Franziska R. Traube, Julian Jude, Philipp Rathert

## Abstract

The CRISPR/Cas9 technology has revolutionized genotype-to-phenotype assignments through large-scale loss-of-function (LOF) screens. However, limitations like editing inefficiencies and unperturbed genes cause significant noise in data collection. To address this, we introduce CRISPR Gene and Transcriptome Engineering (CRISPRgate), which uses two specific sgRNAs to simultaneously repress and cleave the target gene within the same cell, increasing LOF efficiencies and reproducibility. CRISPRgate outperforms conventional CRISPRko, CRISPRi, or CRISPRoff systems in suppressing challenging targets and regulators of cell proliferation. Additionally, it efficiently suppresses modulators of EMT and impairs neuronal differentiation in a human iPSC model. In a multiplexed chromatin-focused phenotypic LOF screen, CRISPRgate exhibits improved depletion efficiency, reduced sgRNA performance variance, and accelerated gene depletion compared to individual CRISPRi or CRISPRko, ensuring consistency in phenotypic effects and identifying more significant gene hits. By combining CRISPRko and CRISPRi, CRISPRgate increases LOF rates without increasing genotoxic stress, facilitating library size reduction for advanced LOF screens.

**Motivation:** The CRISPR technology (CRISPRko/CRISPRi) enables the specific depletion of target genes with fewer off-target effects, facilitating precise investigations of gene function. Despite its benefits, CRISPR applications have limitations. Residual active protein expression mediated by in-frame DNA repair or alternative splicing^1–8^ as well as strong epigenetic regulation and difficulties in sgRNA design to the transcription start site (TSS)^9–12^ hinder the full potential of loss-of-function studies using CRISPRko or CRISPRi. We aimed to achieve robust target gene reduction in order to improve the reproducibility of the CRISPR technology by integrating the widely used CRISPRko and CRISPRi approaches into a single application.

## Introduction

Loss-of-function (LOF) genetic studies are the gold standard for defining direct gene regulatory functions connected to specific cellular phenotypes. LOF approaches based on RNA interference (RNAi), chemical inhibitors, and targeted protein degradation are broadly applicable, but most often lead to the partial suppression of gene function.^13,14^. Previously, the most commonly used LOF approach was RNAi, which is prone to off-target effects and false negatives due to incomplete knockdown of target genes.^13^ In contrast, CRISPR/Cas9 is capable of making highly specific, permanent genetic modifications that are more likely to interfere with the target gene function. The remarkable efficiency and scalability of the CRISPR genome editing approach represent a substantial advancement in LOF studies within mammalian cell culture systems.^15–17^ The most applied CRISPR approaches to date consist of either genetic knockout (CRISPRko) through the introduction of a frameshift mutation, or transcriptional repression using catalytically dead Cas9 (dCas9) fused to an epigenetic effector such as the Krüppel-associated box (KRAB) called CRISPRi.^9,18^ Both methods utilize a 20 nucleotide (nt) guide RNA (sgRNA) to target the Cas9 protein to the desired complementary genomic region carrying the protospacer adjacent motif (PAM).^19^ In the case of Cas9, a DNA double-strand break (DSB) is induced which is preferentially repaired by non-homologous end-joining (NHEJ) introducing insertions or deletions (InDels) thus resulting in frameshift mutations or the introduction of a premature termination codon (PTC) in the respective transcript.^20,21^ For CRISPRi, the dCas9 is recruited to the transcription start site (TSS) of the targeted protein to epigenetically silence its expression.^22^

Besides LOF studies interrogating individual genes, CRISPR represents the ideal genome engineering system for large-scale forward genetic screening approaches to systematically identify new factors that are involved in normal and pathological processes. Such screens have been employed in many studies,^14^ but applications evolving around CRISPRko can show unpredictable outcomes of NHEJ resulting in in-frame DNA repair as well as by alternative splicing^1–8^ leading to residual active protein expression. Without relying on error-prone DNA DSB repair, CRISPRi exhibits a more homogenous response and an improved gene depletion efficiency without inducing genotoxic stress which increases with each DSB leading to severe off-target effects in dual CRISPRko screens.^23^ However, studies have shown that the binding position in the promoter region as well as the native epigenetic landscape play an important role in CRISPRi silencing efficiency, with genes harboring multiple TSSs complicating complete repression.^9–12^ To compensate for possible inefficient CRISPRi/CRISPRko gene suppression, each gene is generally targeted by 5-20 sgRNAs,^11,24,25^ which not only increases the overall variability of the investigated phenotype, but also drastically elevates the costs of library synthesis as well as sequencing depth. This is not ideal for experimental designs in which cell numbers are limited. Nevertheless, such large-scale sgRNA libraries have been deployed to conduct systematic genetic screens to identify essential protein-coding and non-coding genes,^11,18,26–29^ to uncover gene regulatory networks and regulators of disease-associated states^11,30–32^ among others. To date, several strategies have been employed to optimize current CRISPR LOF approaches ranging from dual sgRNA vectors in order to improve on-target and reduce off-target efficiency^33^ or to interrogate genetic interactions,^34^ to the recruitment of additional silencing entities through SUN-tag or MS2-MCP scaffolding^35,36^ and combinatorial recruitment of different transcriptional regulators (LSD1, KRAB) to improve target gene repression^35,37^ with the newest method CRISPRoff, consisting of dCas9 fused to the DNMT3A-3L and KRAB domains.^38^ Moreover, machine learning approaches developed algorithms for predicting improved guide design rules which increased the activity of sgRNA libraries.^11,24,39^ Nonetheless, commonly used libraries target each gene with five or more sgRNAs^14,24^ with the development of newer highly active LOF libraries reducing the amount of sgRNAs to 2 - 4 sgRNAs per gene.^40–42^ The generation of an ultra-compact (1 - 3 sgRNAs per gene), highly active dual CRISPRko or CRISPRi sgRNA library^35,43^ was another recent approach aiming to reduce the constraints imposed by the large size of single guide RNA (sgRNA) libraries and challenges in generating cell models with consistent CRISPRi-mediated knockdown. However, the high representation of genes targeted only with a single construct complicates the robust identification of screen hits and increases the identification of false positive hits due to off-target binding, which adds to the validation burden.^14^ Additionally, in some cases the dual-guide approach was not able to overwrite strong epigenetic marks^35^ and dual-guide libraries targeting only one gene with a single combination are largely ineffective in identifying potential off-target effects. The development of compact, highly active sgRNA libraries would enable CRISPR LOF screens in primary or stem-cell-derived models, *in-vivo* as well as pooled CRISPR screens with spatial transcriptome or proteome resolution with high-content readout and other experimental designs where cell numbers are limiting. To address this problem and to improve the reproducibility of the technology in LOF studies, we have developed a novel CRISPR system, which enables robust target gene reduction through the combination of Cas9 nuclease-mediated DNA cleavage and repressive epigenome editing of the same target gene. Through the fusion of active Cas9 to a powerful transcriptional repressor and the delivery of two sgRNAs from a dual expression construct, this approach makes use of the optimized sgRNAs developed previously for CRISPRko approaches, which already demonstrate high knockout (KO) effects^39^ and simultaneously induces the additional downregulation of residual target transcript expression. Therefore, potential inefficient KO and transcriptional repression resulting from alternative splicing as well as alternative start codons can be addressed through the combinatorial approach preventing the expression of functional truncated target proteins originating from in-frame DNA repair^1,33^. This approach enhances LOF effects, but does not increase reduced cell proliferation due to genotoxic stress caused by DNA damage response, which is observed when inducing DNA cleavage at multiple loci per cell.^23,44^ The system leads to an increase in the overall LOF of the entire cell population allowing for smaller libraries with much lower sgRNA variation per gene and replicate. The combinatorial CRISPR gene and transcriptome engineering (CRISPRgate) approach paired with reduced-scale sgRNA libraries enables systematic forward genetic screens to interrogate the depletion of genes essential for cell growth and the resulting phenotypes with high efficacy and reproducibility.

## Results

### Truncated sgRNAs prevent CRISPR nuclease activity while simultaneously silencing gene expression

To improve CRISPR LOF screens, our approach was directed towards increasing the phenotypic effect by simultaneous gene and transcriptome engineering (CRISPRgate). Repression of target gene expression in parallel to the introduction of a DNA DSB in a shared exon among alternative gene transcripts promised to be the most effective approach (Figure 1A). For this purpose, we fused functional Cas9 to the KRAB domain of ZIM3 (ZIM3), which has previously been reported to show superior silencing efficiency among a large number of KRAB domains.^35,37^ We confirmed these findings, by comparing a fusion of dCas9 with either ZNF10-KRAB or ZIM3-KRAB and subsequent transduction of both fusion proteins into a NIH/3T3 reporter cell line expressing the fluorescent reporter protein mCherry.^45^ Targeting the dCas9 to the promoter, we were able to monitor ZIM3 and ZNF10-KRAB silencing over the course of 14 days and could detect a significantly stronger reduction in reporter gene expression in cells expressing ZIM3-KRAB compared to ZNF10-KRAB (Supplementary figure 1A). To simultaneously achieve strong and continuous transcriptional repression and Cas9-mediated DNA cleavage, the nuclease activity of Cas9 has to be controlled to maintain the recruitment of active Cas9 at the promoter region. It was recently shown that Cas9 DNA cleavage activity is impaired when sgRNAs are shortened from the 5’-end^46,47^ and that targeting of dCas9 can be achieved with 15 nt sgRNAs to activate gene expression.^48^ We tested the ability of truncated sgRNAs to recruit dCas9-ZIM3 to its target region and to induce an efficient and continuous downregulation of gene expression. To this end we applied the same experimental setup as before using the NIH/3T3 cells expressing the *mCherry* reporter construct and dCas9-ZIM3 and transduced these with a set of truncated sgRNAs designed to target the promoter region of the synthetic promoter expressing *mCherry* (Supplementary figure 1B). Recruitment of dCas9-ZIM3 to the synthetic promoter for eight days did not lead to a significant difference in reporter gene silencing efficiency when comparing the 20 nt sgRNA and the PAM-distal truncated sgRNAs (Supplementary figure 1B). Having confirmed that truncated sgRNAs efficiently recruit dCas9 to its target equally affecting transcriptional repression the next step was to determine whether the fusion of Cas9 to ZIM3 and recruitment to the TSS using a truncated sgRNA is different from the commonly used dCas9 using a regular length sgRNA. Therefore, we fused the ZIM3-KRAB domain to the active Cas9 nuclease (ZIM3-Cas9) in a conditional lentiviral expression vector allowing the timed induction of ZIM3-Cas9-expression via Doxycycline (Dox). We transduced either the ZIM3-Cas9 fusion or dCas9-ZIM3 into the erythroleukemia cell line TF-1 and -sorted a cell population expressing moderate amounts of ZIM3-Cas9 or dCas9-ZIM3 by fluorescence-activated cell sorting (FACS). Afterwards, the cells were transduced with an sgRNA-expressing vector targeting two genes encoding non-essential transmembrane receptor proteins. The genes *CD13* (*ANPEP1*) and *CD33* (*SIGLEC3*) are strongly expressed in TF-1 cells and the designed sgRNAs target the TSS of *CD13* and *CD33* using either 20 nt sgRNAs or their respective 15 nt truncations (Figure 1B). The expression of *CD13* and *CD33* was monitored using fluorescently-labeled antibodies against CD13 or CD33. Both the ZIM3-Cas9 with a 15 nt sgRNA as well as dCas9-ZIM3 with a 20nt sgRNA resulted in a reduction of the protein level of CD13 and CD33 with the truncated sgRNA seeming to exhibit a faster gene silencing effect as observed for the 20 nt sgRNA with a significant difference observed after 5 days. After 14 days both sgRNAs exhibit similar endpoint silencing efficiencies (Figure 1B). As a next step, we wanted to investigate the DNA cleavage ability of ZIM3-Cas9 using 15 nt sgRNAs as well as 20 nt sgRNAs and compare this effect to Cas9 alone. Therefore, we transduced the TF-1 cells with Cas9 and sorted cells based on the high Cas9 expression by FACS. Both ZIM3-Cas9 and Cas9 cells were transduced with a vector allowing the dual expression of two sgRNAs expressed from individual promoters (U6 and H1). ZIM3-Cas9 or Cas9 expression was induced through the addition of Dox for 14 days with both 20 nt sgRNAs targeting either *CD13* or *CD33* inducing a strong reduction of CD13 and CD33 in ZIM3-Cas9 as well as in Cas9 expressing cells (Figure 1C). Both ZIM3-Cas9 as well as WT Cas9 had comparable efficiencies with the 20 nt sgRNA guide whereas the 15 nt analog targeting the same genomic region within the gene body of *CD13* failed to induce any loss of functional protein. However, we observe that the 15 nt sgRNA that targets the gene body of *CD33* did induce ∼40% protein reduction in cells expressing ZIM3-Cas9, but not with WT Cas9 (Figure 1C). We terminated the expression of Cas9 and ZIM3-Cas9 through the removal of Dox in cells transduced with sgRNAs where an effect on *CD13/CD33* expression was observed and monitored these cells for the course of 64 days (Supplementary figure 1C). No recovery of *CD13* expression was observed in cells where ZIM3-Cas9 or Cas9 was targeted with the 20 nt sgRNA hinting towards an irreversible DNA DSB induced by ZIM3-Cas9 or Cas9 cleavage (Supplementary figure 1C). The reduction of CD33 observed for the truncated sgRNA when using ZIM3-Cas9 was not stable after the removal of Dox, hinting towards a reversible CRISPRi effect on the 800 bp distant promoter region. In contrast, the identical 20 nt long sgRNA demonstrated an irreversible reduction of CD33, observed after removal of Dox in both ZIM3-Cas9 or Cas9 cells (Supplementary figure 1D). The striking difference observed between both targets when using the truncated sgRNA could be explained by the distance between the targeted region and the TSS. For *CD33* there was a smaller distance between the target and promoter region (∼800 bp) compared to *CD13* (∼2800 bp) (Figure 1C) suggesting that in the case of *CD33*, binding of ZIM3-Cas9 was able to silence *CD33* expression, which was recovered after Dox removal (Supplementary figure 1D). To further validate that the irreversible loss of *CD13* and *CD33* expression observed for the 20 nt sgRNAs was a result of CRISPR-induced DNA cleavage and that the matching 15 nt sgRNA did not induce DNA cleavage we conducted a mismatch-cleavage-assay and observed cleavage products only in cells expressing a 20 nt sgRNA while the 15 nt guide did not depict any CRISPR cleavage products for *CD13* or *CD33* (Supplementary figure 1D). Additionally, we performed amplicon sequencing of the Cas9 cleavage site to quantify the amount of DSB breaks observed for the 15 nt and 20 nt guides (Figure 1D) and did not detect a significantly higher InDel frequency when using the 15 nt guide compared to the control, whereas when using the 20 nt guide a significant increase in InDel frequency was observed. This suggested that in the case of CD33, we observe the simultaneous effect of genome editing and transcriptional interference in the case of the 20 nt sgRNA for the first time demonstrating that our envisioned approach could improve LOF approaches by simultaneous gene editing and transcriptome interference.

**Figure 1:**
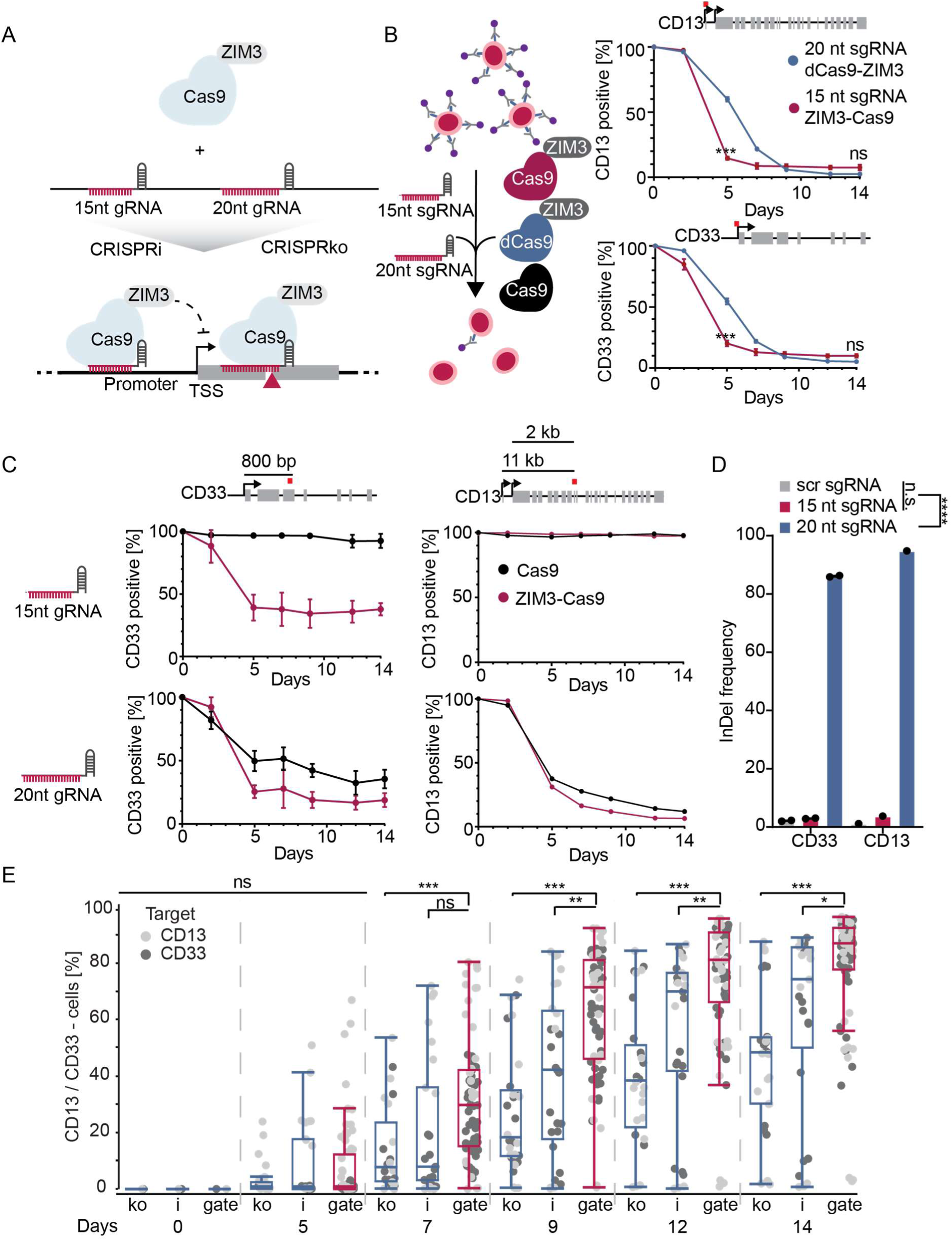
Design and validation of the CRISPRgate concept to improve standard CRISPR LOF approaches. **(A)** Schematic overview of the CRISPRgate setup. An irreversible DNA double-strand break is introduced by targeting ZIM3-Cas9 to an exon of the gene of interest using a 20 nt guide RNA. This is combined with a simultaneous recruitment of ZIM3-Cas9 to the promoter region of the target gene using a truncated guide RNA inducing a stable repression of gene expression. **(B)** Flow-cytometric analysis of the depletion of the non-essential proteins CD13 and CD33 using standard 20 nt long sgRNAs or truncated 15 nt sgRNAs in TF-1 cells expressing the dCas9-ZIM3 fusion protein. (n = 3, mean ± S.D..; n.s. = non-significant, ordinary two-way ANOVA with Tukey post-hoc test). **(C)** Flow-cytometric analysis of the depletion of the non-essential proteins CD13 and CD33 using standard 20 nt long sgRNAs or truncated 15 nt sgRNAs in TF-1 cells expressing either Cas9 or the ZIM3-Cas9 fusion protein. The location of the sgRNA target region is highlighted in red as well as the distance to the nearest TSS (n = 3, mean ± S.D..; n.s. = non-significant, ordinary two-way ANOVA with Tukey post-hoc test). **(D)** InDel frequency determined by deep sequencing of the CD13 and CD33 genomic target region of ZIM3-Cas9 expressing TF-1 cells treated either with Control, 15 nt long or 20 nt long sgRNAs. Statistical analysis was performed on the pool InDel frequency of 15 nt, 20 nt and control samples (****<0.0001, one-way ANOVA) **(E)** Time-resolved quantification of CD13 and CD33 negative TF-1 cells expressing the indicated sgRNAs after induction of ZIM3-Cas9. A total of eight CRISPRko and seven CRISPRi sgRNA designs were combined in 22 CRISPRgate constructs for comparison. Data is displayed as a single datapoint for each sgRNA or sgRNA combination and replicate summarized in a boxplot. (n = 3, mean, box, and whiskers min to max.; **P ≤ 0.01, ***P ≤ 0.001, n.s. = non-significant; two-way ANOVA with a Tukey post-hoc test).

### CRISPRgate significantly accelerates and increases CRISPR-mediated gene depletion

Having validated the core needs for our combinatorial approach, we aimed to investigate if the CRISPRgate system can provide benefits concerning efficiency in the reduction of gene expression and time compared to standard CRISPRi or CRISPRko approaches. To this end, we performed a time-resolved reduction of CD13 and CD33 protein in TF-1 cells using the same inducible ZIM3-Cas9 construct. We designed a set of sgRNAs targeting either the promoter region or the gene body of the target genes and again monitored the reduction of CD13 and CD33 protein by flow cytometry (Figure 1E). In order to investigate a potential combinatorial effect when using CRISPRgate, the *CD13* and *CD33* sgRNAs were chosen based on their predicted *in-silico* depletion efficiencies^49^ (low, medium, and high) and used in combination (CRISPRgate) compared to CRISPRi or CRISPRko alone. Since we found that the 15 nt sgRNA seems to silence gene expression to a similar extent but at a faster pace compared to the 20 nt sgRNA (Figure 1B), we decided to use the truncated sgRNA guides for the CRISPRi comparison to attribute the observed effect to the combination of gene and transcriptome editing rather than a higher silencing pace. Initially, we conducted experiments with three CRISPRgate sgRNA combinations in which sgRNAs with different predicted efficiency were combined. As expected, the combination of two sgRNAs using the CRISPRgate system increased *CD33* suppression (Supplementary figure 2A).

After these supportive initial results, we tested a comprehensive selection of CRISPRko and CRISPRi sgRNAs with a large range of *in-silico* prediction scores,^49^ as well as their respective CRISPRgate combinations for their effect on *CD13* and *CD33* expression over the course of 14 days (Figure 1E). Overall, we observed a significantly stronger effect on *CD13/CD33* expression already after seven days compared to the CRISPRi and CRISPRko controls using the CRISPRgate combinations. This tendency was consolidated over the following days, leading to a significant improvement in the increase of the CD13/CD33 negative cell population ranging from 45% to 95% with almost all CRISPRgate combinations accumulating between 75% - 90% at day 14, whereas only a partial loss of *CD13*/*CD33* expression was achieved with both CRISPRi and CRISPRko sgRNAs (Figure 1E). Strikingly, CRISPRgate not only exhibited the highest silencing efficiency but the overall variance in sgRNA performance of all sgRNAs (percentage CD13 and CD33 negative after 14 days) was also drastically lower for CRISPRgate compared to CRISPRko and CRISPRi (Figure 1E). To assess if the increased depletion efficiencies follow a combinatorial effect we analyzed the amount of CD13/CD33 negative cells at day 9 or day 14 for all CRISPRi sgRNAs with their respective CRISPRko sgRNA combination. Interestingly, we were not able to identify a clear trend for the combinatorial effect (Supplementary figure 2B). However, CRISPRgate is not able to result in a significant increase in gene depletion efficiency when combining two non-functional sgRNAs, whereas the weak effect of a non-functioning sgRNA can be rescued through the combination with a medium or high-performing sgRNA. In this case, CRISPRgate reduces the overall heterogeneity in gene depletion observed in LOF approaches using single sgRNAs and is capable of producing significant gene depletion at a faster rate (Figure 1E and Supplementary figure 2B). Overall, these results demonstrate that the combination of CRISPRko and CRISPRi sgRNAs within the newly designed CRISPRgate system leads to a beneficial LOF effect.

### Truncated and normal-length sgRNAs exhibit similar off-target effects

Due to the dual nature of our approach, we anticipated higher number of off-targets for the truncated 15 nt sgRNA, responsible for transcriptional repression, throughout the genome whereas the 20 nt sgRNA would exhibit similar off-target effects compared to standard Cas9 approaches. Although recent literature highlights that only the first five bases in the seed region influence on- and off-target activity of dCas9 ^50–52^ and therefore the reduction of 5 bp in the prime-distal part should not influence binding, we aimed to determine the transcriptional effects of two sgRNAs targeting *CD33* with several predicted perfect off-targets for the 15 nt sgRNA and compare them to the matching 20 nt sgRNA. The sgRNAs were transduced into dCas9-ZIM3 expressing TF-1 cells and cultivated for 14 days. Afterwards, the cells were harvested and global transcriptome alterations compared to a neutral control sgRNA were determined by RNA-seq. Both the 20 nt sgRNA as well as the 15 nt sgRNA exhibited a similar number of genes with reduced expression compared to their sgRNA counterpart, with most genes having similar expression levels for both sgRNAs (Figure 2A). We next predicted all off-targets of the 15 nt sgRNA with up to 3 mismatches (MM) *in-silico* and filtered for all off-targets which are in the range of - 1000 to 1000 bp to the nearest TSS. We calculated the LFC of the *CD33* targeting sgRNA to the control sgRNA for both 15 nt and 20 nt sgRNAs for these genes and did not detect any significant difference between the 15 nt sgRNA and the 20 nt sgRNA in terms of off-target effect (Figure 2B), which was also evident after investigating the LFC expression of the potential off-targets according to the number of MM (Supplementary figure 3A). Exemplarily we identified a subset of off-targets with a variety of MM, showcasing similar effects of the 15 nt sgRNA and 20 nt sgRNA (Figure 2C). Overall, recruitment of dCas9 to the DNA seems to tolerate more MM than expected, to the extent that there are no significant differences when using a 15 nt sgRNA with 2 MM to the target region or the corresponding 20 nt sgRNA with 4 or 5 MM to the target region (Figure 2C). We validated these findings by targeting dCas9-ZIM3 to a fluorescent reporter system expressing *mCherry* ^45^ using 20 nt sgRNAs and matching 15 nt sgRNAs with a range of different MM (Supplementary figure 3B). Overall, there were no differences in silencing of the *mCherry* expression between 20 or 15 nt long sgRNAs and only an sgRNA with at least two consecutive MM in the seed region impacted the silencing of *mCherry* significantly (Supplementary figure 3B and C). This data implies that off-target prediction and determination performed for Cas9 is not sufficient to predict dCas9 off-target activity where we observe only a mild difference in dCas9 activity with truncated sgRNAs and MM in the seed region, which is in line with current literature.^50–52^

**Figure 2:**
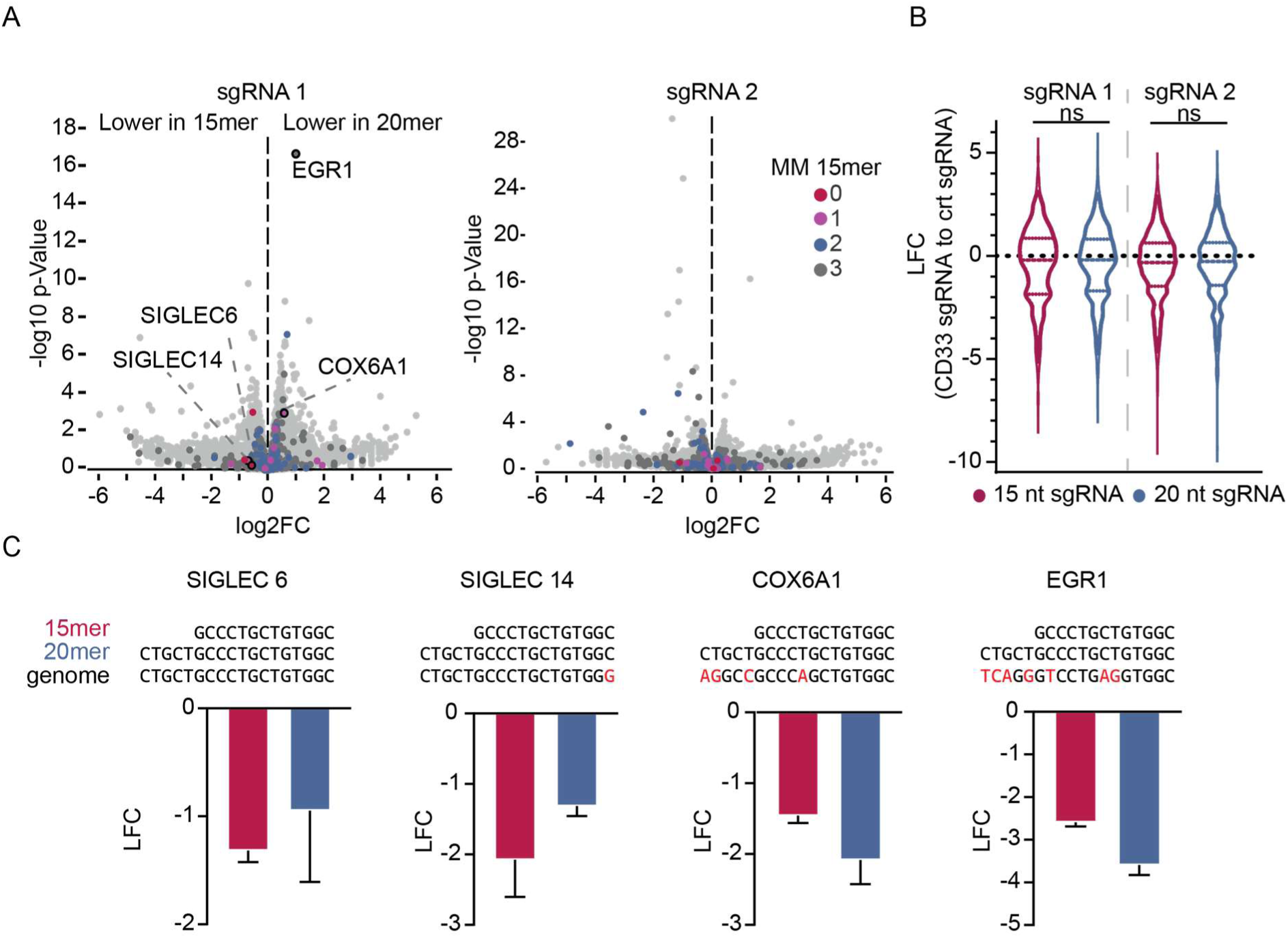
Off-target analysis of 15 and 20 nt long sgRNAs does not reveal significant differences. **(A)** Log2 fold change between identical sgRNAs targeting the TSS of CD33 either 15 nt or 20 nt long. Two different genomic regions within the promoter region of CD33 were chosen as targets. Off-targets towards TSS of other genes were predicted *in-silico* for up to three mismatches for the 15 nt sgRNA and are indicated by color. **(B)** Log2 foldchange of the 15 nt or 20 nt CD33 targeting sgRNA and the sgRNA control for all potential off-targets identified *in-silico* (up to 3 mismatches). **(C)** Log2 foldchange of identified off-targets for the 15 nt sgRNA calculated for both sgRNA variations with the DNA base mismatches indicated in red.

### Assessment of phenotypic consequences following CRISPRgate-mediated BUB1 depletion

Based on the initial results obtained from depletion of *CD13* and *CD33*, we postulated that the enhanced depletion efficiency of CRISPRgate will result in increased phenotypic effects compared to conventional approaches. To evaluate this hypothesis, we wanted to test CRISPRgate on the BUB1 kinase, a crucial protein involved in regulating the progression through the cell cycle as an integral part of the spindle assembly checkpoint (SAC), which is essential for proliferation.^53^ However, the essentiality of BUB1 was under debate^54–57^ since BUB1 KO cells did not exhibit an impaired cell cycle.^56,57^ In line with recent literature^1^ it was shown that BUB1 KO cells still express low levels of functional BUB1 protein.^53,58^ Only the removal of the remaining BUB1 mRNA by RNAi or the complete removal of the *BUB1* gene by CRISPR substantially affected the cell cycle.^59^ Targeting of distinct domains of BUB1 also did not result in a loss of BUB1 functionality, with alternative splicing rendering the indels useless^58,60^ and the generation of full removal of the gene by CRISPR was only successful in haploid cells.^60^ This suggested that BUB1 would be the ideal target to test our CRISPRgate approach since we induce a KO in an early exon and combine this with the simultaneous reduction of the residual mRNA.

We therefore decided to benchmark CRISPRgate against BUB1, which was already deemed as the “zombie” protein in literature,^61^ aiming to assess if the CRISPRgate approach could effectively predict the impact of suppressing BUB1 on the regulation of cell cycle progression and proliferation. Standard LOF screening approaches assess the essentiality of a specific gene by determining the phenotypic effects of 5 to 20 sgRNAs. In order to determine if the increased efficiency observed on the level of non-essential proteins (Figure 1E) can be translated to a potential multiplexed LOF screening approach we aimed to assess if targeting BUB1 would generate comparable phenotypic results targeting a challenging protein, which was shown to express alternative variants after CRISPR/Cas KO.^53,58^ To this end, we selected the top five predicted sgRNAs for CRISPRko and CRISPRi^49^ targeting *BUB1* and sequentially combined the top three sgRNAs to be used in all possible combinations in the CRISPRgate system (Supplementary figure 4A). TF-1 cells expressing conditional ZIM3-Cas9 were transduced with the respective *BUB1* sgRNAs and the fraction of sgRNA-positive cells was monitored over the course of 21 days. Interestingly, although we selected the top 5 predictions for CRISPRi and CRISPRko most sgRNAs were not able to evoke a significant phenotypic effect when compared to the control and exhibited a large variance in the observed negative proliferation effect (Supplementary figure 4A). Strikingly, the combination of the top 3 prediction pairs in the CRISPRgate system led to a significant depletion already observed after seven days, which further increased over the course of three weeks (Figure 3A). Overall, the CRISPRgate sgRNA combinations not only led to a stronger phenotypic effect after a shorter time frame but also displayed less variance resulting in better statistical significance (Figure 3A) suggesting that CRISPRgate achieves significant effects at much earlier time points. With the single CRISPRi sgRNAs targeting BUB1 producing a high heterogeneity in the response to BUB1 depletion, we wanted to assess if transcriptional silencing of BUB1 might be inefficient due to the essentiality of this gene. Therefore, we used the recently published CRISPRoff technology^38^ in combination with three dual sgRNA plasmids targeting the TSS of BUB1. Interestingly, CRISPRoff was not able to reduce the heterogeneous response of transcriptional silencing of BUB1 in TF-1 cells and only resulted in a significant difference compared to the control on day 19 (Supplementary figure 3B and 3C), demonstrating the dependence on a Cas9-induced DSB for a significant BUB1 depletion. To investigate the biological impact of BUB1 depletion on cell cycle progression, we selected *BUB1* sgRNA combinations, which displayed a faster negative proliferative effect, and investigated these in HEK293 cells expressing conditional ZIM3-Cas9 or CRISPRoff (Figure 3B). Additionally, we compared CRISPRgate to conventional Cas9 using a dual sgRNA strategy simultaneously targeting the first and last exon of *BUB1*, which was previously reported to achieve full removal of the *BUB1* gene in HAP1 cells, but not in any other cell line.^60^ Compared to this “benchmark” CRISPRgate showed almost the same negative effect on cell proliferation (Figure 3B), whereas the single CRISPRi and CRISPRko sgRNAs showcased a moderate negative effect on cell proliferation as observed in TF-1 cells (Figure 3A). The effect of BUB1 depletion using CRISPRoff (dual sgRNA) resulted in a faster phenotypic effect compared to CRISPRi, but reached similar end-point values and was not able to increase the negative proliferative effect further (Figure 3B). Interestingly, we observed a gradual decrease of CRISPRoff-positive cells transduced with a scrambled control (scr) over time which we do not observe for CRISPRgate (Supplementary figure 4D) hinting that the expression of CRISPRoff could be toxic through unspecific silencing of other genomic regions. A phenomenon that has already been described several times for DNMT3A-3L constructs.^36,59,62,63^ Additionally, we determined the relative *BUB1* expression of all samples at day 6 using qPCR (Supplementary figure 4E). Overall, CRISPRgate reached the highest depletion efficiency reducing the expression of *BUB1* to 30%, whereas CRISPRi and CRISPRoff reached a reduction of 50%, further highlighting the difficulty of epigenetically silencing tightly regulated genes essential for cell proliferation. To determine whether the negative proliferative effect observed upon *BUB1* suppression is a result of its essential function in cell-cycle regulation, which will result in DNA replication stress and subsequent apoptosis, we stained the described CRISPRgate combinations, CRISPRoff and the respective controls for their DNA content in order to determine different cell cycle stages (Supplementary figure 4F). Both CRISPRgate and CRISPRoff in combination with an sgRNA targeting a non-essential gene did not result in significant differences in cell cycle distribution compared to the HEK WT cells whereas a significantly decreased G1 cell population was observed for all BUB1 depletion samples (CRISPRi, CRISPRko, CRISPRgate, Benchmark and CRISPRoff) (Figure 3C). However, only CRISPRgate and the Benchmark resulted in a significant difference in other cell cycle stages compared to HEK WT. The Benchmark sgRNAs demonstrated a significant accumulation of cells in the S-phase whereas CRISPRgate resulted in a significantly higher cell population in the G2/M phase of the cell cycle, which would be in conjunction with the proposed function of BUB1 in the M-phase of mitosis.^53^ Interestingly, the CRISPRgate sample had an increased cell population in the sub-G1 phase, a cell cycle phase that is an indication of apoptotic cell death, while the sub-G1 population of the CRISPRgate control, the same sgRNAs used for CRISPRi and CRISPRko, the Benchmark sgRNA combination (induction of two DSB) and CRISPRoff samples was hardly noticeable (Figure 3C). These findings imply that the observed phenotype does not stem from higher genome toxicity, but rather from the impaired cell cycle attributed to the improved reduction of BUB1.

**Figure 3:**
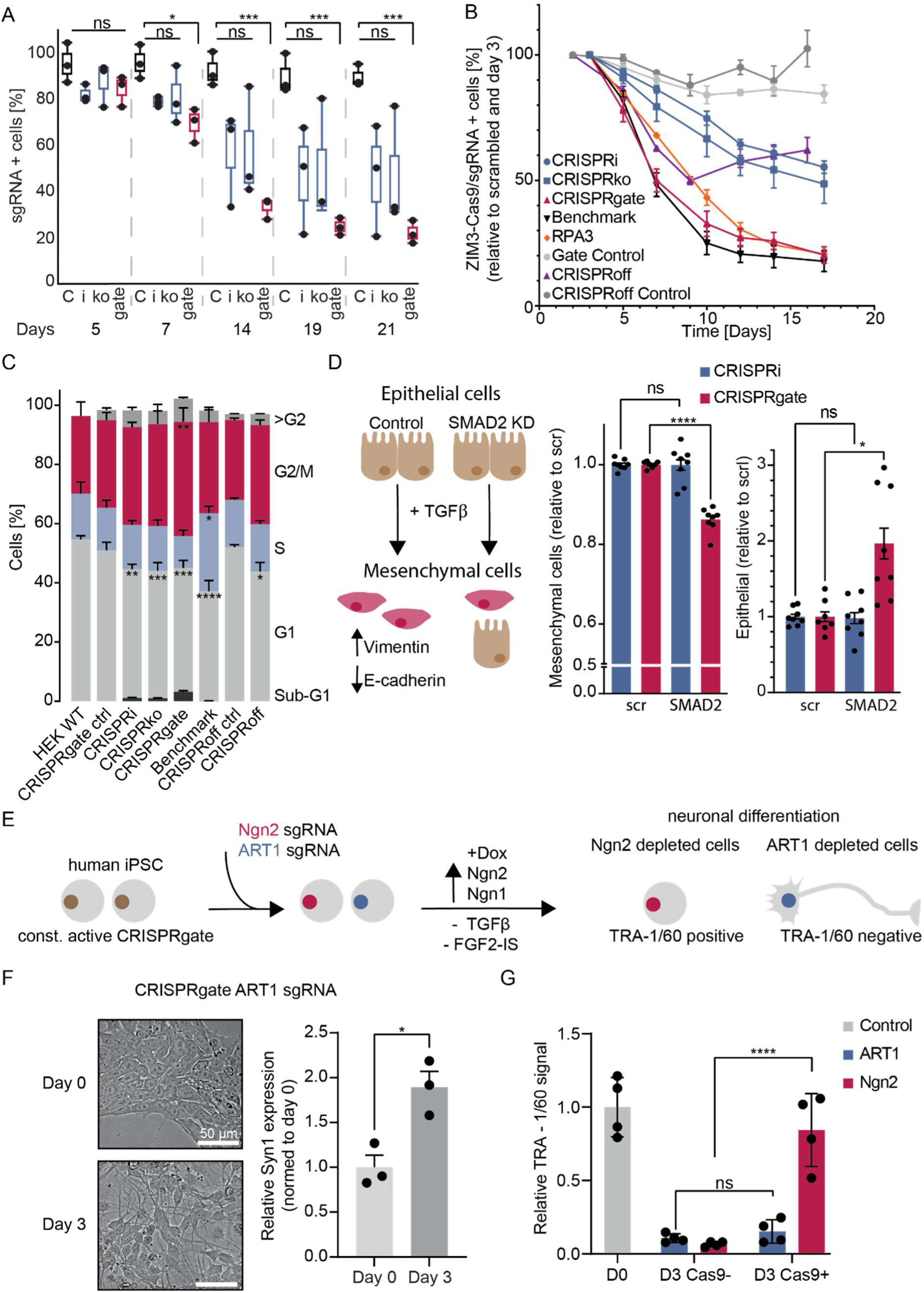
CRISPRgate outperforms standard CRISPR LOF methods and validates the essential role of BUB1 in cell cycle progression. **(A)** Competitive proliferation assays of TF-1 cells expressing the indicated sgRNAs targeting BUB1. Shown is the relative fraction of GFP+/sgRNA+ cells relative to the initial measurement over the course of 21 days (n = 3, mean ± S.D..; *P ≤ 0.05, ***P ≤ 0.001, n.s. = non-significant; two-way ANOVA with a Dunnett post-hoc test). **(B)** Competitive proliferation assays of HEK293 cells expressing ZIM3-Cas9 or the CRISPRoff construct and the indicated sgRNAs targeting BUB1 to validate the improved CRISPRgate effect observed in TF-1 cells. (n = 3, mean ± s.e.m.; *P ≤ 0.05, **P ≤ 0.01, ***P ≤ 0.001, n.s. = non-significant; two-way ANOVA with a Dunnet post-hoc test). **(C)** Inferred distribution of cell-cycle phases of HEK cells harvested at day 5 of BUB1 depletion as indicated in B. Percentages in each phase of the cell cycle were automatically assigned using FlowJo. (n = 3, mean ± s.e.m.; *P ≤ 0.05, **P ≤ 0.01, ****P ≤ 0.0001, n.s. = non-significant; one-way ANOVA with a Bonferroni post-hoc test). **(D)** Epithelial to mesenchymal transition was induced in MCF10A cells expressing either dCas9-ZIM3 or ZIM3-Cas9 and control or SMAD2 targeting sgRNAs using TGFβ. The fold change of cells detected in the epithelial and mesenchymal population was calculated in SMAD2-depleted cells relative to the control (n = 3, mean ± s.e.m.; *P ≤ 0.05, **P ≤ 0.01, ****P ≤ 0.0001, n.s. = non-significant; one-way ANOVA with a Bonferroni post-hoc test). **(E)** Schematic overview of the experimental setup used to validate the tolerability of CRISPRgate in induced pluripotent stem cells. **(F)** Differentiation of iPSCs expressing CRISPRgate and an *ART1* targeting sgRNA was monitored for three days by assessing cell morphology and expression of *Syn1*. **(G)** Relative TRA-1/60 signal (marker for pluripotency) of ZIM3-Cas9 positive and ZIM3-Cas9 negative Dox stimulated iPSCs either expressing an sgRNA targeting *ART1* or the transgene *Ngn2* responsible for neuronal differentiation.

### CRISPRgate extends to non-tumorigenic and stem cell contexts

Next, we wanted to assess if CRISPRgate can be applied in experimental setups involving cells sensitive to DNA damage and epigenetic silencing such as primary cells or stem cells. To test whether we could effectively perturb epithelial to mesenchymal transition (EMT) induced by TGF-β, we transduced the non-tumorigenic epithelial MCF10A cells with dCas9-ZIM3 or ZIM3-Cas9. A cell population was sorted based on the expression of dCas9-ZIM3/ZIM3-Cas9 and displayed normal growth behavior. We then transduced these cells with sgRNAs targeting *SMAD2*, a known mediator of TGF-β signaling as well as a scr sgRNA control and sorted cells based on the expression of the sgRNA. After one week, we induced EMT using TGF-β for eight days and stained for E-cadherin and Vimentin, markers of EMT) (Figure 3D, Supplementary figure 5A). Overall, we did not observe any effects of CRISPRgate cells expressing the scrambled sgRNA on TGF-β signaling dynamics. As expected, cells expressing the scr sgRNA TGF-β showed a reduction of E-cadherin and an increase in Vimentin expression. After *SMAD2* depletion (Supplementary figure 5B) the percentage of cells that underwent EMT was significantly reduced using CRISPRgate (Figure 3D). In contrast, no significant shift was observed when using CRISPRi targeting *SMAD2* (Figure 3D). Importantly, the cells that did not react to TGF-β-induced EMT due to downregulation of *SMAD2* by CRISPRgate did not exhibit negative effects on viability due to genotoxic stress. Instead, they remained in the epithelial cell population (Figure 3D).

As a next step, we validated CRISPRgate in settings involving more sensitive cells such as induced pluripotent stem cells (iPSCs). To this end, we transduced constitutive active ZIM3-Cas9 into human iPSCs which were modified to activate neuronal stem cell differentiation upon Dox induction through the expression of the mouse transgenes *Neurogenin-2* (*Ngn2*) and *Neurogenin-1* (*Ngn1*) connected via a P2A element (Figure 3E).^64^ We designed CRISPRgate sgRNA combinations, one targeting a non-essential gene expressed in neuronal cells (*ART1*) and another sgRNA combination targeting the transgene *Ngn2*. We selected the cells expressing the sgRNA using G418 for seven days, validated the loss of *ART1* via qPCR (Supplementary figure 5C), and induced neuronal differentiation. Overall, the cells expressing the *ART1* targeting sgRNA showed the anticipated morphological changes after three days (Figure 3F) and the upregulation of *SYN1* (Figure 3F), a gene that is activated in neuronal differentiation.^64^ Three days after induction, the cells were harvested and stained for a cell surface pluripotency marker (TRA-1/60), which is reduced after differentiation to neuronal cells.^64^ We measured a significant loss of the TRA-1/60 signal compared to the uninduced WT, without finding any significant differences between the ZIM3-Cas9 negative and ZIM3-Cas9 positive cells expressing the sgRNA targeting *ART1* (Figure 3G). This indicates that the iPSCs have normal phenotypic behavior and signaling which is not impaired through potential DNA damage stress by the expression of CRISPRgate. In order to prove that CRISPRgate is able to interfere with the cellular differentiation, we analyzed TRA-1/60 expression after Dox induction in the cells, where the *Ngn2* transgene was targeted. Whereas ZIM3-Cas9 negative cells showed the same reduction of Tra-1/60 as the *ART-1* targeted cells, ZIM3-Cas9 positive cells showed only a minor reduction with a significantly higher relative TRA-1/60 expression compared to the ZIM3-Cas9 negative cells (Figure 3G), indicating that neuronal differentiation was successfully impaired by CRISPRgate.

### CRISPRgate improves LOF screen performance

Having illustrated that our CRISPRgate approach improves current CRISPRi/CRISPRko in terms of depletion efficiency and performance, which resulted in comparable phenotypic effects for almost all sgRNA combinations and less overall variance in the proliferative effect (Figure 1E and Figure 3A, B), we postulated that screening using CRISPRgate would be able to evoke a significant phenotypic effect at a much faster rate due to the enhanced gene suppression efficiency combined with less sgRNA per gene variance leading to improved hit calling at reduced library coverage. We therefore decided to test the CRISPRgate system using a multiplexed LOF phenotypic screening approach. To this end, we collaborated with Twist Bioscience to design and clone oligo fragments consisting of a 15 nt sgRNA and its tracr followed by a promoter sequence and the 20 nt sgRNA (Supplementary figure 6A). In order to synthesize the dual sgRNA on a single <300 nt long oligonucleotide fragment, we made use of a minimal engineered hybrid H1 promoter in which all core elements except for the Staf-domain were substituted with elements from the 7SK promoter (minH1/7SK).^65^ This minimal promoter construct was reported to show only Pol III lacking any Pol II activity,^65^ and we did not identify any significant differences in the effect on *CD33* suppression between the H1 and minH1/7SK promoters over the course of 14 days (Supplementary figure 6B). Interestingly, the activity of the hU6 promoter was significantly enhanced by the exchange of the H1 to the minH1/7SK leading to a faster and stronger reduction in *CD13* expression in the tested experimental setup (Supplementary figure 6B). Using the minH1/7SK promoter we synthesized and cloned a CRISPRgate oligonucleotide pool designed to target 1137 genes (including 10 internal controls) involved in chromatin regulation.^66,67^ For each gene, three independent CRISPRgate combinations were designed, combining the top three CRISPRi (shortened to 15 nt) and CRISPRko sgRNAs, which were selected based on the most recent prediction algorithms and designs. ^39,49^ For genes with multiple annotated transcription start sites (TSS) with a distance of >1000 bp from each other, the same CRISPRko sgRNAs were combined with different 15 nt sgRNAs targeting each TSS (Supplementary figure 6C). The initial library pool consisting of 270 nt oligonucleotides was synthesized and quality-controlled using deep sequencing after cloning into the final screening vector and the overall library distribution was calculated (Supplementary figure 6D). The CRISPRgate library harboring 3686 sgRNAs in total was transduced in triplicate into three independent conditional ZIM3-Cas9 TF-1 cell clones. After selection with Neomycin (NEO) for 7 days, ZIM3-Cas9 expression was induced by the addition of Dox (Figure 3A). Based on the encouraging effects observed with suppression of *BUB1* (Figure 3A and B), where a negative proliferative effect was observed at much earlier time points and with much higher reproducibility between different sgRNA combinations when using CRISPRgate compared to CRISPRi or CRISPRko, cells were passaged only 7 times (14 days, 70 h doubling time). We determined the chimera rate, which arises due to different steps during the screen, such as homologous recombination during library amplification in *E. coli*, a template-switch mediated by lentiviral reverse transcriptase, or PCR template swapping during Illumina library preparation. This can produce a recombined element with two sgRNAs targeting different genes.^35^ Overall, we started the screen with an initial chimera rate in the oligo pool of 9.4% which increased after library preparation ranging from 14% to 20% per single cell clone (Supplementary figure 7A). Library preparation using the pooled dual sgRNA library resulted in a chimera rate ranging from 34.61% to 38.03%, which was most likely a result of unfavorable PCR conditions, which were optimized to amplification of sgRNA sequences from genomic DNA leading to conditions favoring template swapping in the plasmid pool. After sequencing, we filtered for reads where the 15 nt sgRNA and the 20 nt sgRNA matched the designed oligonucleotide targeting the same gene. Overall, the CRISPRgate screening approach achieved a noteworthy reproducibility between all three replicates (Supplementary figures 8 and 9) and we determined the performance for all sgRNAs using the CRISPRBetaBinomial or CB^2^ algorithm.^68^ To assess the screen performance, we calculated the depletion efficiency of selected positive and neutral control sgRNAs. Neutral control sgRNAs targeted non-essential *CD13/CD33* genes whereas positive controls were directed against *RPA3*, *MYC*, and *CSF2RA*, a receptor subunit of the granulocyte colony-stimulating factor (GCSF), which is essential for TF-1 proliferation (Supplementary figure 7B). As expected neutral control sgRNAs did not show a strong effect on cell proliferation in contrast to positive sgRNAs (Figure 4B). Next, we investigated the effect of individual sgRNAs targeting a set of established common essential and non-essential genes.^69^ Targeting essential genes led to negative LFC values contrary to sgRNAs targeting non-essential genes, which clustered around LFC values from 0 to 1. This effect was also visible on the gene LFC level (Figure 4C and Supplementary figure 7C). To evaluate the performance, we determined the area under the curve (AUC) of the ROC curve calculated by using all sgRNAs targeting the gold-standard established common essential and non-essential genes present in the CRISPRgate library (Supplementary figure 10A), which demonstrated that the CRISPRgate approach, with a substantially reduced number of sgRNAs per gene and passages, can compete with many previously published high-end CRISPR library screening approaches(Supplementary figure 10A).^11,24,29,35,39–41,43^ We calculated the Spearman correlation between sample replicates for read counts, sgRNA and gene LFC and observed a high reproducibility throughout the replicates for the CRISPRgate screening approach on all layers, showcasing that our approach enables a high consistency in performance and can compete with previous methods (Figure 4D and Supplementary figure 8 and 9). Our previous data suggested that CRISPRgate reduces the overall noise within phenotypic effects among sgRNAs targeting the same gene (Figures 1 and 3). To confirm this observation, we compared the ΔLFC (maximum LFC-minimum LFC) between sgRNAs targeting the same gene to the identical genes in recently published LOF screens.^11,24,29,35,39–41,43^ Considering only the sgRNAs targeting essential genes, we noticed a significantly lower ΔLFC for sgRNAs targeting the same gene in the CRISPRgate screen compared to other screening approaches where three or more sgRNAs per gene were used (Figure 4E). Additionally, this was also the case when comparing all genes targeted, indicating an overall more consistent performance for sgRNAs targeting the same gene than observed in other screening approaches (Figure 4E). A similar trend was observed when we investigated the variance of the LFC for all sgRNAs targeting the same gene (Supplementary figure 10B). To evaluate if the reduced variance and highly consistent performance of the sgRNAs targeting the same gene is reflected by an improved hit calling, we compared the −log10 of the adjusted p-value (FDR) for each sgRNA in published screens filtered for genes present in the CRISPRgate chromatin library (Figure 4E). The analysis showed that sgRNAs exhibited a much higher reproducibility in significant phenotypic effects in the CRISPRgate screen compared to the other analyzed public screens (Figure 4E and Supplementary figure 11), which was also reflected by the gene LFC range (Figure 4D). Interestingly, the high significance observed with some of the published LOF screens across replicates on sgRNA level was not conferred to the gene level suggesting that the large heterogeneity observed when using multiple sgRNAs to target the same gene increases the variance leading to a reduction in significance on gene-LFC. With the CRISPRgate approach, this loss in significance was not observed (Figure 4E). As a result of the low ΔLFC between sgRNAs and reduced variance observed in the CRISPRgate screen, the low p-values per sgRNA are transferred to the overall adjusted p-value per gene. This leads to more consistent gene hits with a very high significance with the CRISPRgate screening approach.

**Figure 4:**
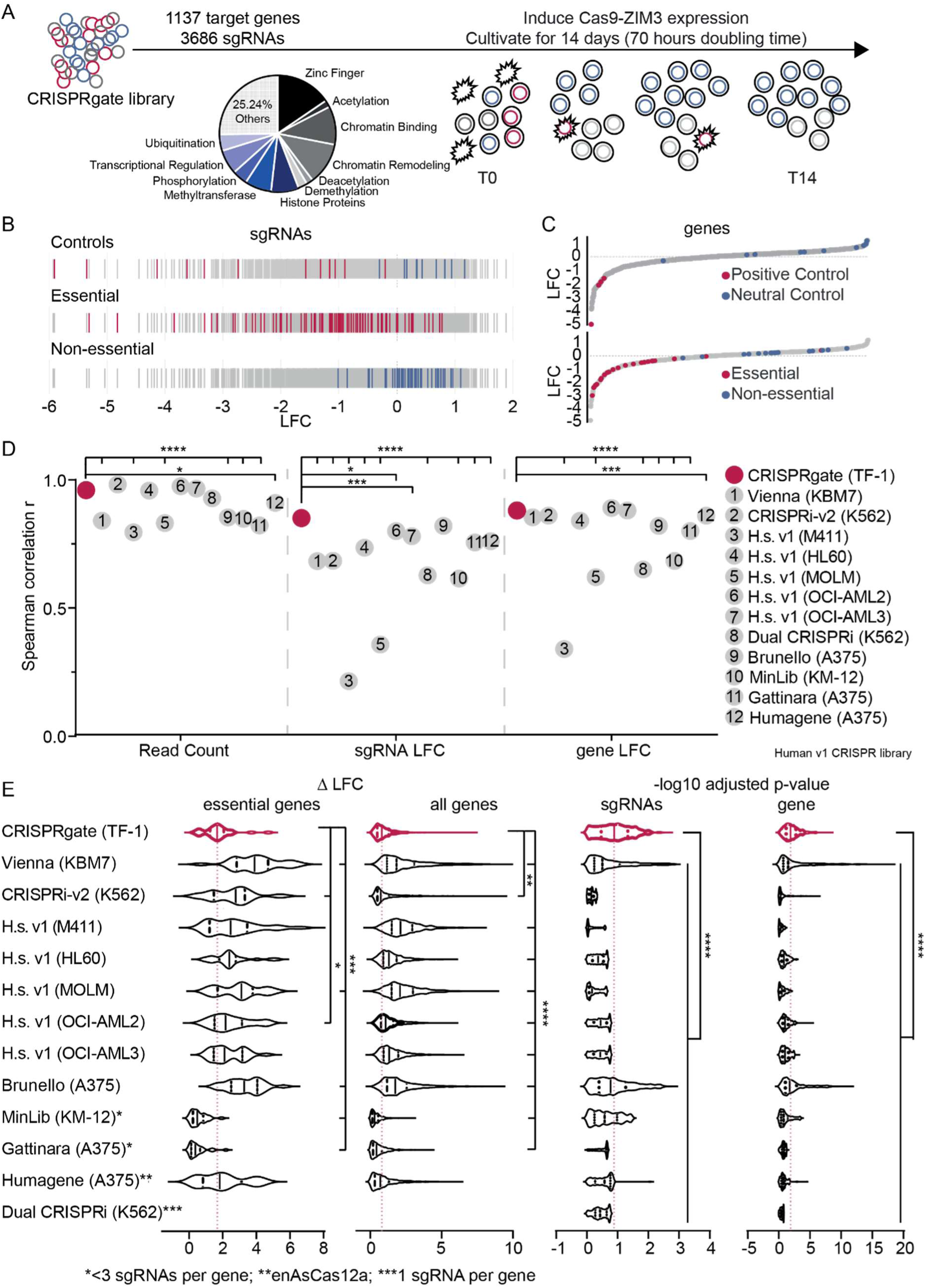
CRISPRgate outperforms published LOF dropout screen revealing an improved sgRNA consistency. **(A)** Schematic overview of the screening setup used for the CRISPRgate screen. A library composed of 3686 sgRNAs targeting 1137 chromatin-related genes was virally transduced into TF-1 erythroleukemia cells expressing ZIM3-Cas9. After antibiotic selection, cells were treated with DOX for 14 days. **(B)** Performance of individual sgRNAs. The Log fold change (LFC) of all sgRNAs targeting internal positive (red) and neutral controls (blue) as well as sgRNAs targeting known essential (red) and non-essential genes (blue) is depicted. Data are averaged across 3 individual replicates. **(C)** Scatter plot depicting all genes ranked by the average LFC of all sgRNAs per gene across all three replicates. Internal positive (red) and neutral (blue) control genes (top), as well as essential (red) and non-essential genes (blue) are highlighted (bottom). **(D)** Spearman correlation r calculated for the replicates of the CRISPRgate screen and replicates of published screen datasets on read count, sgRNA LFC and sgRNA LFC level. **(E)** Violin plots comparing the CRISPRgate system with a set of published CRISPR screening approaches. Left: comparison of the ΔLFC (maximum LFC – minimum LFC) of sgRNAs targeting the same gene depicted for essential genes and all genes investigated in the screen. Right: comparison of the -log 10 adjusted p-value distribution at the sgRNA and gene level. The black vertical lines depict the median for each screen and the red dashed line is the median of the CRISPRgate screen. (*P ≤ 0.05, ***P ≤ 0.001, ****P ≤ 0.0001; one-way ANOVA with a Dunnett post-hoc test).

### CRISPRgate generates improved dropout effects through a combinatorial effect of CRISPRi and CRISPRko

To determine if the improved performance of the CRISPRgate library (Figure 4) was due to a combination of a CRISPRko and CRISPRi sgRNA targeting the same gene in one construct, as suggested by our initial experiments (Figures 1 and 3), we compared sgRNA dropout effects of genes where two different TSSs were targeted. Indeed, we found a set of genes where we observed a strong LFC difference when targeting the different TSS (Supplementary figure 12A), which did not correlate to the distance between both TSS (Supplementary figure 12B). Additionally, to rule out a possible epigenetic silencing effect originating from a CRISPRko sgRNA targeting near the TSS, we plotted the LFC of sgRNAs targeting essential genes against the relative distance of the CRISPRko sgRNA to the TSS and did not identify any correlation (Supplementary figure 12C) suggesting that we do not observe epigenetic silencing over long distance through retention of ZIM3-Cas9 after cleavage. We then determined the functional TSS (F-TSS) and non-functional TSS (NF-TSS) of the affected genes, which revealed a highly significant difference in gene depletion upon targeting a different TSS (Figure 5A and Supplementary figure 12D). For instance, CRISPRgate sgRNAs targeting the strongly selective genes *ADAR* and *ERG* (DepMap),^70,71^ resulted in a relatively moderate negative LFC when sgRNAs were directed against the NF-TSS. However, combining the same CRISPRko sgRNAs with sgRNAs targeting the F-TSS led to a drastic LFC decrease with all three CRISPRko sgRNAs (Figure 5B). Since the CRISPRi sgRNAs used to assemble the CRISPRgate constructs were adapted from the CRISPRi-v2 library,^11^ we were able to compare the effects of targeting individual TSSs of the essential gene *DDX46*^69^ in both CRISPRi-v2 as well as CRISPRgate screens. The effect observed with sgRNAs for the F-TSS resulted in a strong negative LFC in both screens, only the CRISPRgate approach was able to produce a negative LFC when targeting the NF-TSS (Figure 5C). This improved performance was also visible when comparing false positive rates determined by non-essential genes and plotted against the true positive rate for both screens (Figure 5D) indicating that the combination of CRISPRi with the CRISPRko leads to a significantly improved gene dropout effect in multiplexed screening approaches (Supplementary figure 12E).

**Figure 5:**
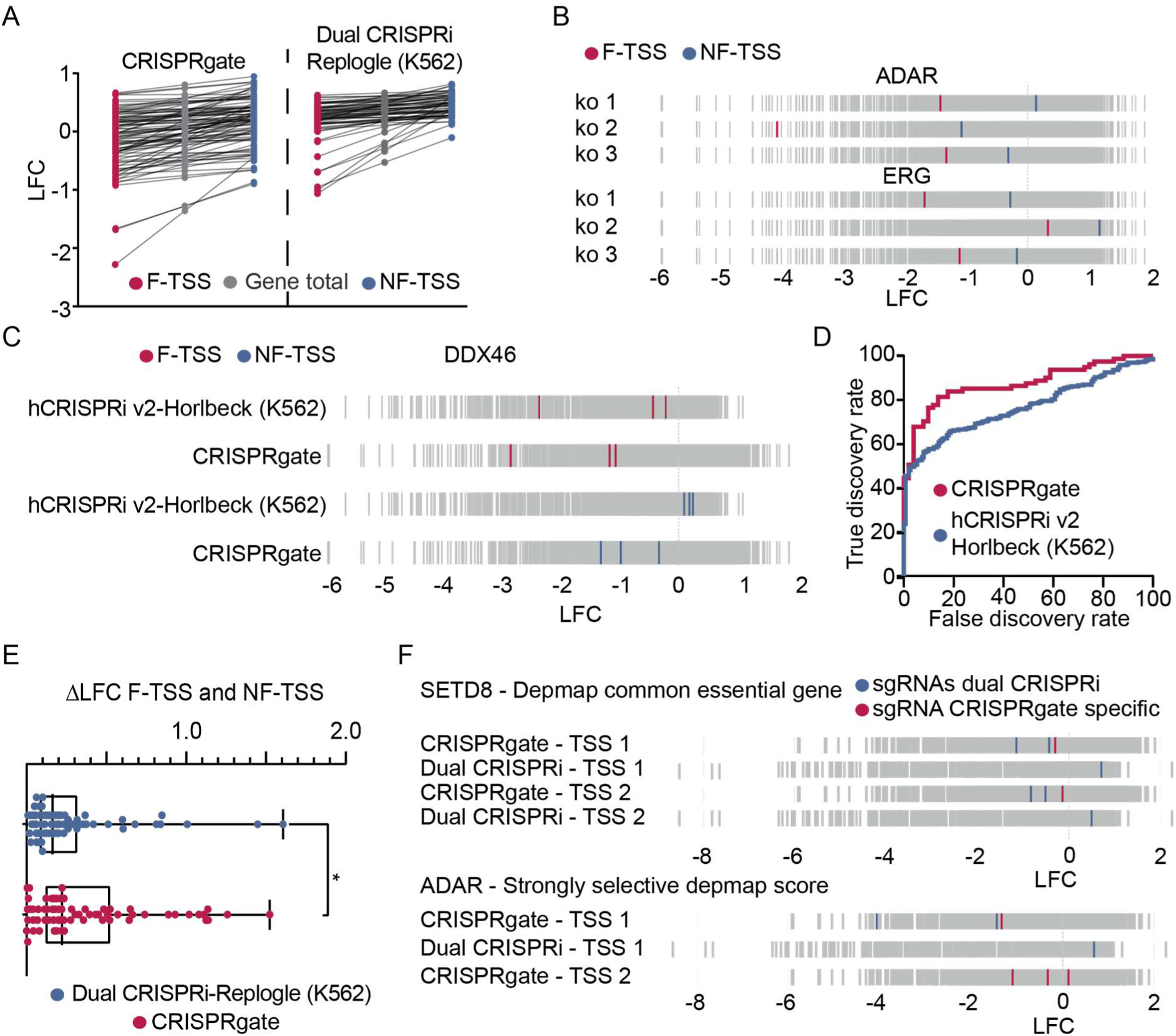
Combination of CRISPRi and CRISPRko has a highly additive effect in a multiplexed LOF screening setup. **(A)** Box plot comparing the depletion of genes where more than one TSS was targeted with different sgRNAs highlighting the differential depletion effect comparing functional (F) and non-functional TSS (NF) in the CRISPRgate screening approach to the effect observed in a published screening dataset using the same sgRNAs (28). **(B)** Performance of individual sgRNA combinations demonstrating that the CRISPRgate improves the KO effect observed when targeting the NF-TSS compared to when targeting the F-TSS of essential genes **(C)** Individual sgRNA performance, exemplified for the essential gene DDX46, indicating that the sgRNA combination of CRISPRgate rescues nonfunctioning CRISPRi indicated by the depletion observed when the NF-TSS is targeted compared to a published CRISPRi screen using the same sgRNAs (16). **(D)** ROC sensitivity curve of the CRISPRgate compared to a published CRISPRi screen (16) based on sgRNAs targeting essential and non-essential genes. **(E)** Individual sgRNA performance comparison of CRISPRgate with a published screening approach employing a dual CRISPRi sgRNA strategy targeting the same TSS (28). The CRISPRi sgRNAs identical in both screens are highlighted (blue) as well as the CRISPRi sgRNA solely used in the CRISPRgate screen (red). **(F)** Box plot depicting the ΔLFC difference when targeting the F and NF TSS comparing CRISPRgate with a published dual CRISPRi screening dataset (28). (*P ≤ 0.05; non-parametric Wilcoxon test).

In a recently published dual sgRNA CRISPRi library, dual sgRNA constructs were designed to either target the same TSS or to target two different TSSs^35^ based on the same CRISPRi-v2 library, which we used to devise the CRISPRgate library.^11^ To verify the combinatorial phenotypic effect of our CRISPRgate library, we conducted the same analysis for the dual CRISPRi screen and observed a significantly higher ΔLFC between the sgRNAs targeting individual TSSs of the same gene in the CRISPRgate approach compared to the dual CRISPRi screen (Figure 5E). The combination of the dual CRISPRi sgRNAs with a CRISPRko sgRNA in the CRISPRgate setup improved the dropout effect (Figure 5F) illustrated for *SETD8*, a DepMap common essential gene, as well as for *ADAR,* a DepMap-annotated strongly selective gene (Figure 5F). These data suggest that the additional CRISPRko effect has a higher influence on the LFC compared to targeting the same TSS with two CRISPRi sgRNAs. This was also apparent when comparing the individual performance of sgRNAs targeting essential genes in the CRISPRgate TF-1 screen with LOF screens performed with the CRISPRi-v2 library and dual CRISPRi library (Supplementary figure 12E). The CRISPRgate approach showed an improved performance compared to the CRISPRi-v2 library in terms of efficiency and consistency (Figure 5D), indicating that our approach of combining a CRISPRi sgRNA with a CRISPRko sgRNA is a considerable addition to the toolbox of so far published screening methods.

### Hit validation illustrates specific CRISPRgate benefits

Finally, we decided to benchmark the CRISPRgate screen to the dependency score derived from the DepMap portal (https://depmap.org/portal/) for the same cell line.^70,71^ First, we compared the essential genes targeted in the CRISPRgate library, identifying similar effects in the CRISPRgate screen and the DepMap annotation (Supplementary figure 12F) where we observed a comparable performance. After K-means clustering we observed 4 distinct groups (Figure 6A). The first group is composed of genes that show no antiproliferative effect (LFC and DepMap score > 0) and the second group demonstrates a negative proliferative effect in both screens. The third group is comprised of genes that exhibit a strong negative DepMap score being either positive or slightly negative in the CRISPRgate screen and the fourth group represents genes with a strong negative LFC detected in the CRISPRgate screen but having a positive or slight negative DepMap score (Figure 6A). In total, the last cluster is comprised of 147 genes of which we chose *TDRD12*, *KDM1A*, *GFI1B*, and *DPF1*, which show a very minor depletion score (DepMap) and a strong phenotypic effect in the CRISPRgate screen and aimed to validate the CRISPRgate benefits in single competitive proliferation assays (Figure 6D). To account for a potential false-positive phenotypic effect, we assessed the performance of all sgRNAs targeting the respective genes, confirming that all single CRISPRgate sgRNAs resulted in a negative LFC (Figure 6C). Interestingly, *KDM1A* (*LSD1*) was recently shown to exhibit residual mRNA and protein levels in CRISPRko approaches due to alternative splice sites, which rescue the functional impact of mutations^1–8^ and could explain the weak phenotypic effects observed in the previous screens^70,71^ although TF-1 cells are sensitive to LSD1 inhibition.^72^ In addition, GFI1B was shown to play an important role in LSD1 biology^73^ and scored as the top 20 hit in our screen, whereas it exhibited only a weak DepMap score (Figure 6B). We compared the phenotypic effects of our CRISPRgate constructs to the individual sgRNAs used to design the CRISPR gate construct (CRISPRi-v2 and Vienna Bioactivity CRISPR score)^11,39^ as well as the Brunello/AVANA sgRNAs^24^ used to derive the dependency score from DepMap. In all three cases, CRISPRgate outperformed the single sgRNA approaches (Figure 6D) confirming the high potential of the CRISPRgate approach to deliver conclusive results in multiplexed screening approaches, which were not detected by other screening methods previously. So far, the central role of the LSD1 and GFI1B interaction in leukemia^72,73^ has never been observed in a multiplexed LOF screening approach although LSD1 activity has been shown to be essential in TF-1 cells and the LSD1-GFI1 regulatory axis is critical for the proliferation.^72,73^ Interestingly, the respective CRISPRko sgRNAs targeting *GFI1B* or *KDM1A* had a significantly reduced effect on cell proliferation with the CRISPRi in the CRISPRgate combination contributing the most towards the negative proliferative effect. In contrast, targeting *DPF1* or *TDRD12* in single validation experiments, a higher effect on cell proliferation was observed for CRISPRgate and CRISPRko, whereas CRISPRi was unable to induce a negative effect on cell proliferation. This data further highlights the beneficial effect achieved by the combination of CRISPRi and CRISPRko demonstrating the novelty and relevance of the CRISPRgate system for future LOF approaches.

**Figure 6:**
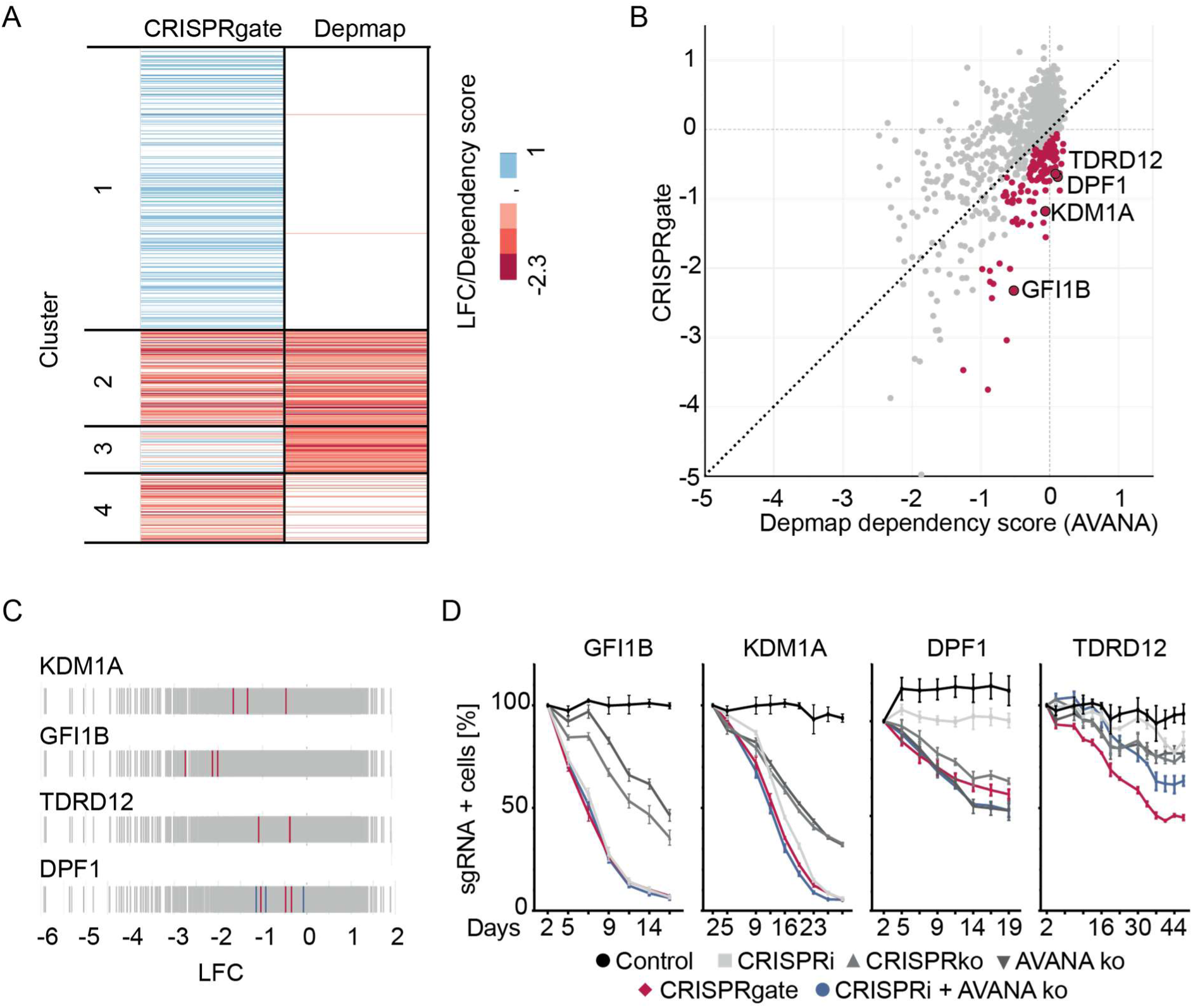
CRISPRgate can identify novel dependencies in multiplexed LOF screening approaches. **(A)** Heatmap showing the LFC of genes targeted in the CRISPRgate screen compared to the dependency score of TF-1 cells from DepMap and clustered into four distinct groups using k-mean clustering. **(B)** Scatter plot showing the CRISPRgate gene level depletion compared to the dependency score from the DepMap portal (https://depmap.org/portal/) for TF-1 cells. Cluster 4 is depicted in red and the differential genes selected for validation of the improved CRISPRgate effect are highlighted with a black circle. **(C)** Individual sgRNA performance of sgRNAs targeting the indicated differential genes. **(D)** Competitive proliferation assays in TF-1 cells expressing ZIM3-Cas9 and the indicated sgRNAs. For the validation, the highest performing CRISPRgate combination and the resulting single sgRNA controls as well as the best *in-silico* predicted CRISPRko sgRNA from the Avana library (19) used to contribute to the DepMap dependency score and a CRISPRgate combination using this KO sgRNA was used. (n = 3, mean ± s.e.m.).

## Discussion

Pooled LOF genetic screening approaches are highly effective in investigating specific phenotypes like proliferation or cell survival. In recent years, the utilization of Cas9 resulted in the replacement of RNAi methods as the gold standard for multiplexed phenotypic screens. To account for sgRNA performance heterogeneity, 5-20 sgRNAs per gene are used in many library pools increasing the size and synthesis costs of the pool as well as the overall variance of the observed phenotypic effect for each gene. Processing genome-wide screens requires large numbers of cells, and is time-consuming and costly, especially in the context of high-content read-out single-cell CRISPR sequencing screens. This can be addressed through library size reduction by screening gene sets comprising several thousand genes, by improving *in-silico* sgRNA prediction, or by designing libraries in which each gene is targeted with a minimal set of sgRNAs in order to achieve a more cost-effective high-throughput screening setup. However, the application of less than three sgRNAs will lead to the discovery of false positive hits (Supplementary figure 12E). Therefore, the development of improved tools leading to reduced library size is highly desired. Many improvements were introduced at the level of the sgRNA design using *in-silico* sgRNA prediction tools to identify efficient sgRNAs to assemble powerful sgRNA libraries.^11,24,39^ Improved performance of these pooled sgRNA libraries resulted in ventures such as the Achilles project performing LOF screens investigating the effects of genes on cellular proliferation and survival over a broad range of cells resulting in gene dependency scores.^25,74,75^ Nevertheless, problems associated with in-frame DNA repair, alternative splice sites, and additional start codons downstream of the original ATG still require the application of more than 3-10 sgRNAs per gene for proper statistics and hit-calling. Similarly, CRISPRi-based LOF screening approaches suffer from limitations leading to ineffective suppression of gene expression,^18,37,76,77^ which was slightly improved by the application of different CRISPRi effector proteins.^18,35,37,76,77^ Previous efforts using machine learning to develop guide design rules increased the activity of sgRNA libraries,^11,24^ but commonly applied libraries still target each gene with more than 3 sgRNAs in order to decrease false-negative screen hits. This hinders a broader application of pooled CRISPR LOF screening approaches with complex molecular readouts or when cell numbers are limited.

To address these problems, we implemented a combinatorial approach combining CRISPRko and CRISPRi utilizing a set of two distinct sgRNAs to introduce a KO at the gene level and simultaneously interfere with residual gene expression of the same gene. To achieve this, we used a normal 20 nt long sgRNA to target functional Cas9 fused to ZIM3 to a common exon of the target gene while providing a truncated 15 nt long sgRNAs to target the same construct to the TSS of the target gene (Figure 1). Our initial results demonstrated a significant benefit in combining both, CRISPRi and CRISPRko, directly resulting in an increased significant phenotypic effect (Figure 1 and Figure 3). We validated that CRISPRgate is also able to be applied in non-tumorigenic cells such as MCF10A or iPSC. Simultaneous KO and repression of a non-essential gene did not impair stem cell proliferation and differentiation signaling dynamics (Figure 3). We confirmed that CRISPRgate can also be used as a LOF tool for genetic dropout screens (Figure 4) and proved that CRISPRgate performs similarly to comparable pooled LOF screens resulting in a phenotypic effect even when the individual CRISPRi or CRISPRko sgRNAs failed to induce a negative proliferative effect (Figure 5 and Figure 6).

A core part of the CRISPRgate approach is the truncation of the sgRNA targeting the TSS of the target gene to 15 nt, not inducing Cas9 DNA cleavage, which could lead to an immediate release of Cas9 binding at the TSS in case the PAM site is lost during NHEJ. In theory, the reduction in sgRNA length would result in an increase in perfect off-targets compared to a standard 20 nt long sgRNA. However, previous reports investigating Cas9 off-target activity determined the first 8-10 PAM-proximal bases to have the strongest effect towards on-target affinity of Cas9, with single base MM within this region severely affecting Cas9 affinity to the target DNA,^46,78,79^ whereas multiple mismatches in the PAM-distal part of the sgRNA are well tolerated and still lead to the induction of a DNA DSB.^46^ A large portion of Cas9 off-target analysis involves DNA cleavage assays rather than biochemical recruitment assays of Cas9 to DNA. More recent studies investigating dCas9 off-target binding also confirm that mismatches within the seed region led to severely reduced dCas9 association rates^50,80^ with biochemical recruitment analysis further demonstrating that the binding of Cas9 only depends on a few bases (minimum 5 bases) within the seed region.^51,52^ Our data investigating the off-target effects of two truncated sgRNAs and their 20 nt analogs (Figure 2 and Supplementary figure 3) supports these findings showing that the last 5 bp are not essential for off-target activity and we did not identify a significant difference between the truncated and normal-length sgRNA indicating that Cas9 tolerates MM more than anticipated in cases when DNA cleavage is not needed. To account for this, we designed three CRISPRgate sgRNA combinations per gene. In order to be considered as a true hit at least two of the three sgRNAs need to show a comparable trend. Libraries with fewer sgRNAs per gene would be unable to identify these off-targets efficiently (Supplementary figure 12G).^50^ Due to the high reproducibility observed for different sgRNAs targeting the same gene the library size can be reduced substantially while maintaining the opportunity to call hits based on statistical significance which would make CRISPRgate a valuable addition to the current CRISPR toolbox to perform LOF screens.

Dual sgRNA libraries require special caution in library design, monitoring of the chimera rate after synthesis, cloning as well as increased costs due to longer DNA synthesis required for cloning dual sgRNA constructs. By utilizing the minimal H1/7SK promoter we were able to reduce the dual sgRNA construct <300 bp, allowing to synthesize high-quality, highly accurate oligo pools leading to reduced errors and a low chimera rate (Supplementary figure 7A) compared to other dual libraries, which report ∼30% chimera rate.^35^ However, several sophisticated cloning strategies for the generation of dual-sgRNA libraries have already been published, which can be employed. Instead of synthesizing a large oligo harboring both the sgRNAs as well as the minimal promoter and tracr employed for the CRISPRgate library cloning (Supplementary figure 6A) a 90 bp oligo can be used to generate a dual sgRNA library reducing the overall synthesis costs.^34,35,81–83^ However, additional cloning steps could lead to an increase in sgRNA chimeras due to homologous recombination in *E. coli* during library amplification, sgRNA recombination between lentiviruses as well as template swapping during sequencing preparation leading to a recombined dual sgRNA cassette with sgRNAs targeting different genes.^11,84,85^ To monitor this problem and to be able to filter false sgRNA combinations we adapted our sequencing strategy towards paired-end sequencing, which delivered the information for correct sgRNA pairing for each construct.

Overall, our initial findings that the combination of gene and transcriptome engineering improves individual CRISPRko and CRISPRi LOF approaches were confirmed in a multiplexed screening approach, ranking comparable to or higher than other published screening approaches in terms of consistency in performance across sgRNAs targeting the same gene, thereby increasing the significance of hit calling. Strikingly, we were also able to prove a combinatorial effect of CRISPRi sgRNAs improving ineffective KO sgRNAs and vice versa showcasing CRISPRgate to provide a valuable complement to existing CRISPRi and CRISPRko LOF screening approaches. Finally, we compared our screening results to the DepMap portal and identified a cluster of novel genes (147) which when depleted resulted in a negative phenotypic effect not previously identified in DepMap of which we validated 4 hits. However, we also noted a cluster of genes with a negative DepMap score (99) and only mild effects in the CRISPRgate screen. Interestingly further analysis revealed that 17% of the genes in this cluster are histone proteins and their variants, which are known to have a low turnover rate^86^ and would have therefore needed more passages to evoke negative proliferative effects, which were applied in the screen deposited in the DepMap portal ^71^.

Based on our results, we envision that CRISPRgate could enable genetic screening for complex phenotypes with large-scale sgRNA libraries in experimental settings where cell numbers are limited or small sgRNA libraries are required. Our setup not only allows for combinatorial transcriptional repression and KO approach, but can also be used to screen dual CRISPRi or CRISPRko libraries in LOF dropout or orthogonal screening approaches. This powerful CRISPR screening workflow could enable the identification of novel dependencies in *in-vivo* mouse or PDX models and primary patient samples left uncovered by *in-vitro* screens. Combined with emerging technologies such as single-cell and time-resolved transcriptomics,^87–89^ orthogonal screening approaches^90^ or organoid model screening,^91^ CRISPRgate could prove to be a novel tool, which enables the characterization of genetic perturbations on the global transcriptome at single-cell resolution and in settings with limited required cell numbers.

### Limitations of the study

CRISPRgate is a combination of both CRISPRi and CRISPRko and cannot fully rescue the target gene reduction effect when both sgRNAs are non-effective. This problem can be minimized by selection of >3 sgRNA designs per target gene and more sophisticated sgRNA design algorithms. In addition, we observed high chimera rates our study occurring during library amplification for sequencing, which could also be occur through template swapping and homologous recombination. As discussed above, this is a main factor complicating the use of dual sgRNA libraries, which can be controlled through the development of more sophisticated library cloning approaches and optimized PCR conditions. Furthermore, CRISPRgate employs a truncated sgRNA to deliver ZIM3-Cas9 to the respective target gene in order to induce gene repression without the introduction of a DSB. In theory this would lead to a significant increase in of perfect off-targets throughout the genome. However, our data confirms other studies showing that Cas9/dCas9 targeting is only weakly affected by bases in the PAM-distal part of the sgRNA and we observe that the 20 nt sgRNA has a similar number of off-targets as the corresponding 15 nt.

## ACKNOWLEDGEMENT

We thank all laboratory members for constructive discussions throughout the project and Regina Philipp as well as Ama Amoateng for technical support. We are grateful to A. Jeltsch for advice throughout this study. We thank the Twist Bioscience team in particular Tavneet Gill, Xianan Liu, Caitlin Hoeber, Carlo Antonio Bilbao for cloning the CRISPRgate Library. pCAG-Eco was a gift from Arthur Nienhuis & Patrick Salmon. pCMVR8.74 was a gift from Didier Trono.

We thank Johannes Zuber (The Research Institute of Molecular Pathology (IMP), Vienna, Austria) for sharing reagents; and all members of the Rathert and Jeltsch labs for reagents, protocols, and discussions.

The work described here was supported by the Wilhelm-Sander Foundation (2016.082.1, 2020.055.1) and the German Cancer Aid (70113426). The work of S.A.J and Z.N was funded by the Robert Bosch Stiftung. The work of C.B. and F.R.T. was funded by CRC1309 (Grant Nr. 325871075, Project C08).

## Author contributions

J.J., and P.R. co-developed the idea. J.S. planned and performed experiments and analyzed data. J.W. cloned the Cas9 constructs. J.S., J.J., and P.R. designed the library. C.B., L.H., H.M., M.K., S.L., and F.R.T. contributed critical reagents, performed and analyzed experiments. Z.N. and S.A.J. contributed critical reagents and performed Illumina sequencing. S.L., F.R.T., S.A.J., and J.J. provided critical advice and support. P.R. designed experiments, analyzed data, and supervised this research. J.S. and P.R. co-wrote the paper. All authors have read and agreed to the published version of the manuscript.

## Supplemental information titles and legends

### Figures S1-S12

Supplementary Tables S1-6.xlsx: Supplementary Table 1: Primers; Supplementary Table 2: CRISPRgate Library sgRNAs; Supplementary Table 3: Normalized screen count of all analyzed screens; Supplementary Table 4: sgRNA guide sequence; Supplementary Table 5: Normalized RNA-seq count and all DE-seq analysis; Supplementary Table 6: Amplicon sequencing count

## STAR★Methods

### Resource availability

#### Lead contact

Further information and requests for resources and reagents should be directed to and will be fulfilled by the lead contact, Philipp Rathert.

#### Materials availability

Plasmids generated in this study will be deposited to Addgene upon publication.

#### Data and code availability

- The raw CRISPRgate screening files, the off-target RNAseq as well as amplicon-seq of the truncated and non-truncated sgRNAs are available at GEO and will be publicly available as of the date of publication.
- This paper does not report original code.
- Any additional information required to reanalyze the data reported in this paper is available from the lead contact upon request.

### Key resources table

| REAGENT or RESOURCE | SOURCE | IDENTIFIER |
| --- | --- | --- |
| Antibodies |  |  |
| Anti-CD90.1(Thy1.1)-APC | Thermo Fisher | AB_469420 |
| Anti-CD90.1(Thy1.1)-APC | BioLegend | AB_1595470 |
| PE anti-human CD33 | BioLegend | AB_314347 |
| PE anti-human CD13 | BioLegend | AB_314179 |
| anti-CD324 (E-Cadherin) - clone 4A2 monoclonal mouse | Cell signaling | AB_2728770 |
| anti-Vimentin - clone D21H3 - monoclonal rabbit | Cell signaling | AB_10695149 |
| Alexa Fluor 647 goat anti-mouse IgG (H+L) | Thermo Fisher | AB_2536165 |
| Alexa Fluor 488 goat anti-rabbit IgG (H+L) | Thermo Fisher | AB_143165 |
| TRA-1-60 Antibody, anti-human, REAfinity™ | Miltenyi Biotec | AB_2654228 |
| Bacterial and virus strains |  |  |
| *E. coli* Stbl3™ | Invitrogen | C737303 |
| Chemicals, peptides, and recombinant proteins |  |  |
| Recombinant human GM-CSF | Miltenyi Biotec | 130-095-372 |
| hHOLO-Transferrin | Merck Millipore | 616424 |
| hrInsulin | Sigma-Aldrich | I9278 |
| hFGF-2-IS | Miltenyi Biotec | 130-104-921 |
| hrTGF-β1 | Miltenyi Biotec | 130-095-067 |
| Deposited data |  |  |
| CRISPRgate screen, RNA-seq and amplicon seq | This paper | https://www.ncbi.nlm.nih.gov/geo/query/acc.cgi?acc=GSE238225 |
| Experimental models: Cell lines |  |  |
| NIH3T3 | ATCC | CRL-1658 |
| LentiX293T | Takara Bio | 632180 |
| HEK293T | DSMZ | ACC 635 |
| TF-1 | ATCC | CRL-2003 |
| iNGNs | Collaboration partner | N/A |
| MCF10A | ATCC | CRL-10317 |
| Oligonucleotides |  |  |
| Primers for cloning, amplification cleavage site, qPCR and illumine sequencing see Table S1 | This paper | N/A |
| CRISPRgate library see Table S2 | This paper | N/A |
| sgRNAs used in this study see Table S4 | This paper | N/A |
| Recombinant DNA |  |  |
| Plasmids will be deposited upon publication |  |  |
| Software and algorithms |  |  |
| Graphpad Prism 5 | Graphpad Software, Inc | N/A |
| Illustrator CS6 | Adobe | N/A |
| Excel 2016 | Microsoft | N/A |
| Tableau 2023.2 | Tableau | N/A |
| R | The R Project | N/A |
| R-studio | Posit | N/A |
| Other |  |  |
| Normalized screen count of all analyzed screens in this publication see Table S3 | This paper | N/A |
| Normalized RNA-seq count and DE-seq analysis see Table S5 | This paper | N/A |
| Amplicon sequencing count see Table S6 | This paper | N/A |

## EXPERIMENTAL MODEL AND STUDY PARTICIPANT DETAILS

### Cell Culture

All media were supplemented with 10% Fetal Bovine Serum, 4 mM L-Glutamine, 10 mM HEPES, 1 mM Sodium pyruvate solution, 100 U/mL Penicillin and 100 µg/mL Streptomycin. NIH/3T3 (male), Lenti-X293T (female) and HEK293T (female) cells were cultivated in DMEM high glucose media (Sigma-Aldrich) and TF-1 (male) cells were cultivated in RPMI 1640 supplemented with 2 ng/mL of recombinant human GM-CSF (130-095-372 Miltenyi Biotec). MCF10A cells were seeded in in DMEM/F12 medium (Thermo Fisher, #21331020) supplemented with 100 ng/ml cholera toxin (Sigma, #C8052), 20 ng/ml epidermal growth factor (EGF) (Preprotech, #AF-100-15), 10 ug/ml insulin (Sigma, #I9278), 500 ng/ml hydrocortisone (Sigma, #H0888), GlutaMax (Thermo Fisher, #35050038), 5 % horse serum (Thermo Fisher, #16050122), 1x penicillin/streptomycin (Thermo Fisher, #15140122). iNGNs ^64^ were grown on Geltrex (ThermoScientific #A1413202)-coated tissue plates and cultivated at 37 °C in water-saturated, CO_2_-enriched (5%) atmosphere. Uninduced iNGNs were cultured in hiPSCs-medium (1:1 DMEM:F-12, GlutaMAX-Supplement (Gibco #10565018), supplemented with 0.2 mM L-ascorbic acid 2-phosphate, 77.6 nM sodium selenite, 10.90 mM NaCl, 0.1 mM nicotinamide, 10 µg/mL hHOLO-Transferrin (Merck Millipore #616424), 20 µg/mL hrInsulin (Sigma-Aldrich #I9278), 20 ng/mL hFGF-2-IS (Miltenyi Biotec #130-104-921), 2.0 ng/mL hrTGF-b1 (Miltenyi Biotec #130-095-067).

## Method details

### Plasmids

The DNA sequence encoding ZIM3-KRAB (ZIM3), without the stop codon, was synthesized and cloned in front of Cas9 in a Dox inducible pRRL-TRE3G-Cas9-P2A-GFP plasmid kindly provided by the Zuber lab ^30^ using standard cloning methods. ZIM3-Cas9 expression was coupled to GFP via a P2A element. Additionally, antibiotic resistance against Blasticidin driven by a PGK promoter was added to select for positively transduced cells (TRE3G-ZIM3-Cas9-NLS-P2A-GFP-PGK-BlastR). For the constitutive CRISPRgate vector, the ZIM3 element without the stop codon was cloned in front of Cas9 of an EF1as-Cas9-P2A-GFP plasmid using standard cloning methods. Additionally, a Blasticidin resistance driven by a PGK promoter was added. For dual sgRNA expression, a dual filler plasmid was cloned adding an additional tracr and promoter to a lentiviral single sgRNA expression vector kindly provided by the Zuber lab ^30^ using a hU6 promoter to express the first sgRNA and either the H1 or minimal H17SK to express the second plasmid. Successful transduction of the dual sgRNA expressing plasmid was monitored via an outer membrane protein (Thy1.1) driven by an Ef1a short promoter and coupled to Neomycin resistance via a P2A element (hU6-filler1-tracr-H1/minH17SK-filler2-tracr-Ef1as-Thy1.1-P2A-Neo).

### sgRNA and library cloning

The CRISPRi and CRISPRko sgRNAs were ordered as complementary oligos harboring overhangs fitting to the BsmBI restriction site of the respective fillers. The complementary sgRNA oligos were phosphorylated, annealed and afterwards cloned into the dual-sgRNA filler plasmid using golden-gate assembly.

For the pooled sgRNA library we designed 270 bp long oligo fragments consisting of the 15 nt CRISPRi sgRNA, a tracr, the minimal H1/7SK promoter and the 20 nt CRISPRko sgRNA. TWIST Biosciences pool synthesized the oligo fragments and cloned these into the filler of a PRRL lentiviral sgRNA vector (PRRL-PBS-hU6-filler-tracr-Ef1as-Thy1.1-P2A-Neo) harboring a primer binding site for PCR amplification for Illumina sequencing kindly provided by the Zuber lab ^30^. TWIST performed library quality control and deep sequencing, identifying an initial chimera rate of 9.78%.

### Cell culture, lentiviral transduction, generation of Tet-On competent cells and single-cell clones

Generation of Tet-on competent cells was performed as previously described ^66^. For lentiviral packaging of pRRL-vectors, plasmids were mixed with helper plasmids pCMVR8.74 (Addgene plasmid #22036) and pCAG-Eco (Addgene plasmid #35617) and 3x (w/w) excess of polyethyleneimine 25K in DMEM. The mix was added dropwise to LentiX cells at 70 - 80% confluency. Media was exchanged after 12 and 24 h. The virus particles were harvested 48h after transfection with an optional second and third harvest after 56 to 72 h. To prevent contamination of the target cells with LentiX, the virus was filtered using a 45 µm filter. pRRL vectors for expression of ZIM3-Cas9 were introduced into the Tet-on competent target cells by transduction at a transduction efficiency <20% to ensure single plasmid integration. For selection, TF-1 cells were treated with 4 µg/mL Blasticidin. The dual sgRNA expression vector was transduced and the cells were selected with 500 µg/mL G418 solution for 7 days. Afterwards, expression of ZIM3-Cas9 was induced by the addition of 1 µg/mL Doxycycline (Dox). Successful integration and expression of the desired plasmids was analyzed two days after transduction and monitored throughout selection and flow cytometry using a MACSQuant Vyb flow cytometer. To generate single-cell clones for the CRISPRgate screen, expression of ZIM3-Cas9 was induced in TF-1 cells and single-cell clones were sorted based on the GFP fluorescence using the Sony SH800S FACS. The single-cell clones were monitored based on the GFP signal, keeping the cells that were able to reversibly induce ZIM3-Cas9 expression by addition and removal of Dox. We then validated the single-cell clones by transducing these with sgRNAs targeting CD13 and CD33, induced the expression of ZIM3-Cas9 using Dox and monitored the depletion of CD13/CD33. We chose three distinct sgRNAs which all showed similar CD13/CD33 depletion efficiencies and were able to specifically respond to Dox induction and removal.

### Immunodetection of CD13 and CD33 depletion

sgRNAs targeting CD13 and CD33 (Supplementary Table 4) were stably integrated using lentiviral transduction. ZIM3-Cas9 expression was induced by the addition of 1 µg/mL Doxycycline (Dox) and the loss of functional CD13 and CD33 was monitored every two days for 14 days through AB staining of CD13 and CD33 and detection by flow cytometry.

### RNA-seq for off-target detection

Two sgRNAs targeting *CD33*, were selected based on their *in-silico* predicted off-target activity (minimum of one perfect off-target for the 15mer) and cloned as standard 20 nt sgRNA or the respective 15 nt truncation. The sgRNAs targeting *CD33* as well as a scrambled sgRNA (scr) were transduced into dCas9-tagBFP-ZIM3 expressing cells and selected for one week. After two weeks of sgRNA expression, the cells were sorted based on the dCas9-ZIM3 and sgRNA expression, harvested and RNA was isolated using the RNeasy Plus Mini Kit (QIAGEN, Hilden, Germany). RNA concentration and quality were assessed using the 260/280 and 260/230 ratios obtained at the NanoDrop. We enriched for mRNA using the NEBNext® Poly(A) mRNA Magnetic Isolation Module (NEB) and used this as input for library generation using the NEBNext® Ultra™ II RNA Library Prep (NEB). The quality and concentration of the cDNA library was assessed using the BioAnalyzer 2100 high sensitivity DNA kit. In case adapter fragments were detected, size exclusion DNA purification using NEBNext® Sample Purification Beads (NEB) was performed according to the manufacturer’s instructions. The samples were sent for deep sequencing and the obtained reads were filtered using Trimmomatic ^92^ removing adapter contamination and filtering reads based on sequencing quality, keeping high-quality reads for further analysis. Next, we aligned both forward and reverse reads to the human genome (hg38) using HISAT2 ^93^ and obtained the gene count files, which were used for DESeq2 analysis ^94^ comparing both sgRNAs (15 vs 20) with each other as well as comparing the individual sgRNAs and the scr control.

### Competitive proliferation assay

TF-1 cells were transduced with the indicated sgRNAs (Supplementary Table 4) and ZIM3-Cas9 expression was induced using Dox. The amount of Cas9 and sgRNA positive cells were measured after two days and set as reference. The proliferation of ZIM3-Cas9 and sgRNA-positive cells was monitored for the indicated time points. HEK293T cells were transduced with the indicated sgRNAs and selected for one week. Afterwards, 80% of sgRNA-positive cells were mixed with 20% WT cells, and expression of ZIM3-Cas9 was induced using Dox. Two days after, the amount of sgRNA and ZIM3-Cas9 positive cells were measured and set as reference, and the proliferation was monitored for the indicated time points.

### Cell cycle analysis

200 000 HEK293T cells were harvested after five days of BUB1 depletion and fixed in 70% ice-cold ethanol for 2h. Afterwards, ethanol was removed by centrifugation and the pellet was resuspended in 200 µL cell cycle staining solution (100 µg/mL RNAseA, 50 µg/mL PI, 0.1% TritonX in PBS) and incubated for 30 to 60 minutes at room temperature in the dark. Directly after incubation, the DNA content was measured by flow cytometry and the cell cycle tool (model Dean-Jett-Fox) of FlowJo was used to automatically detect the cell cycle phase distribution of the different samples.

### Gene expression analysis

Cells were harvested after five days of BUB1 depletion, and mRNA was extracted by using the RNeasy Plus Mini Kit (QIAGEN, Hilden, Germany). RNA concentration and quality was assessed using the 260/280 and 260/230 ratios obtained at the NanoDrop. Reverse transcription was performed with 500 ng of purified RNA using oligo(dT)_18_ primers for the Multiscribe Reverse Transcriptase (Invitrogen) according to the manufacturer’s instructions. Quantitative PCR reactions were carried out using the CFX Connect Real-Time System from Bio Rad using the ORA SEE qPCR Green ROX H Mix (highQu) and the human *BUB1* and *ACTIN* primer set (Supplementary Table 1). The following cycling conditions were performed: 95°C for 3 min, 39 cycles of 95°c for 5 s then 60°C for 30s with the cycling conditions for the melt curve performed afterwards.

### Amplicon sequencing and T7 endonuclease assay

TF-1 cells were transduced with 20nt or 15nt long sgRNAs targeting either CD13 or CD33 as well as with a scr control sgRNA and selected for seven days. ZIM3-Cas9 expression was turned on using Dox and the cells were harvested after 14 days. Primers were designed to amplify the sgRNA targeting region either for amplicon sequencing or mismatch cleavage assay (Supplementary Table 1). Genomic DNA of > 1 Mio sgRNA positive and WT cells was amplified and extracted from an agarose gel and the DNA concentration was determined. For amplicon sequencing, Illumina adapters were added to the PCR product using a second PCR step, the resulting PCR amplicons were then extracted from an agarose gel. The size, concentration, and purity of the DNA amplicon was assessed using the BioAnalyzer 2100 high sensitivity DNA kit. All amplicons were pooled and sent for Illumina sequencing. For analysis, the reads were split based on the internal barcode, the site of cleavage was identified and the percentage of InDels was calculated for the scr control, 15 nt and 20 nt sgRNAs. For the T7 assay, 200 ng of PCR product (WT and sgRNA) was stepwise annealed and T7 endonuclease was added. Cleaved fragments were determined using agarose gel electrophoresis.

### TGFβ induced EMT

MCF10A cells stably expressing dCas9-ZIM3 or ZIM3-Cas9 were transduced with sgRNAs targeting SMAD2, the cells were stained using PE-labeled antibodies against Thy1.1 (receptor co-expressed with the sgRNA) and sgRNA-positive cells were sorted two days after transduction. After one week of sgRNA expression cells were supplemented with 1:1000 Doxycycline to induce the expression of ZIM3-Cas9. One day after initial seeding, MCF10A cells were treated for 8 days with 100 pM TGFβ1 (Prepotech, #100-21C) with re-stimulation every 48 h, and harvested by TrypLETM (ThermoFisher, #12604013). For cell fixation and permeabilization, 100.000 cells per condition were treated with 4 % PFA in DPBS for 15 min, and 0.1 % Triton in FACS media (DPBS + 5 % FHS) for 15 min at 4 °C. Cells were then incubated with anti-CD324 (E-Cadherin) antibody (1:150) and anti-Vimentin antibody (1:150) for 1 h. After blocking in FACS media for 10 min, cells were incubated with secondary antibodies Alexa Fluor 647 goat anti-mouse IgG (H+L), (1:1000) and Alexa Fluor 488 goat anti-rabbit IgG (H+L) (1:250) for 45 min. All antibody incubations were performed in FACS media. Cells were washed between fixation with DPBS and during antibody staining twice with FACS media and afterwards analyzed by flow cytometry. For validation of the SMAD2 depletion, a subset of cells was harvested before TGFβ stimulation, and the RNA was isolated and reverse-transcribed into cDNA. Using specific primers for *SMAD2* (Supplementary Table 1) the relative expression compared to *ACTIN* was measured using qPCR as described in “Gene expression analysis”.

### Neuronal stem cell differentiation and analysis of Tra-1/60

iNGNs were passaged in the uninduced state when they reached about 70 – 80% confluency. For passaging, 2 µM (final concentration) of thiazovivin (Merck Millipore 420220) was added to the hiPSCs medium for 24 h, afterwards medium was changed to hiPSCs-medium without thiazovivin. iNGNs were stably transduced with EF1a-ZIM3-Cas9-P2A-GFP-PGK-BlastR and selected for one week using 2µg/mL blasticidin. Successful selection and expression of ZIM3-Cas9 was monitored by measuring the expression of GFP at the MacsQuant. Afterwards, sgRNAs targeting ART1 or Ngn2 were stably transduced and cells were selected for one week using 100 – 200 µg/mL G418 solution. For induction of neuronal differentiation, the medium was changed to induction medium (hiPSCs-medium without FGF2-IS and TGF-β1), and Doxycycline (Sigma-Aldrich #D9891) was added to a final concentration of 0.5 µg/mL.

### Multiplexed CRISPRgate LOF screen

A CRISPRgate library designed to target 1137 target genes involved in chromatin regulation using a set of 3686 sgRNAs (Supplementary Table 2) was transduced into three independent TF-1 cell clones with comparable ZIM3-Cas9 expression levels of with an sgRNA representation of 1 000 X. After 7 days of antibiotic selection, ZIM3-Cas9 expression was induced using Dox and cells were cultivated for 14 days (7 passages, 70 h doubling time). During cultivation, sgRNA representation and ZIM3-Cas9 expression was monitored using flow cytometry. After 14 days (7 passages), genomic DNA for the three single-cell clones was isolated using phenol-extraction using PhaseLock tubes, followed by ethanol precipitation. Multiple parallel 50 µL PCR reactions, each containing 1 µL gDNA template adding up to 40 µg gDNA and 300 ng for the library pool, using the AmpliTaq Gold Polymerase (Life Technologies) were performed to maintain sgRNA representation. In a first round of PCRs, random barcodes and sample barcodes were added to the sgRNA sequences using the following cycling parameters: 95°C for 10 min; 28 cycles of (95°C for 30 s, 54°C for 45 s and 72°C for 60 s); 72°C for 7 min. PCR products for each single cell clone were combined and purified using the NucleoSpin Gel and PCR clean-up kit (Macherey-Nagel). Afterward using a second round of PCR, using similar cycling conditions with the exception of 10 ng input as template and 7 total cycles the standard Illumina P7 and P5 adaptors were added. All primers used for the library preparation are listed in Supplementary Table 1. The final libraries were cleaned up from a 2% agarose gel, pooled, and analyzed on a P2 flow cell (400 mio reads) using the Illumina NextSeq 2000 with a 35% PhiX spike-in (75 bp paired-end), using standard Illumina primers. Sequence processing was performed using a custom Galaxy workflow (www.usegalaxy.eu). sgRNA count data as well as data used for comparison from other screens are provided in Supplementary Table 3. Forward and reverse reads were combined and non-mapped reads as well as sgRNA chimeras were filtered out. The read count for each sgRNA combination was normalized as count per million using the r-studio tool “CB2” ^68^. Using the same tool fold depletion of individual sgRNA combinations as well as for the targeted genes was calculated.

### Statistical Analysis

All details regarding statistical analysis are provided the respective figure legends and figures, including numbers of replicates for each experiment, statistical tests used and the obtained p values. Results are presented as means ± standard error of the mean [s.e.m.]. If not stated otherwise, statistical significance was calculated by two-way ANOVA with a post-hoc test indicated for each experiment with p≤0.05 considered statistically significant. Statistical significance levels are denoted as follows: ****P ≤ 0.0001; ***P ≤ 0.001; **P≤ 0.01; *P ≤ 0.05; n.s. = non-significant.

