## Supplementary figures for "CRISPR gene and transcriptome engineering (CRISPRgate) improves loss-of-function genetic screening approaches"

### Supplementary Material

Supplementary Figure 1: Validation of the CRISPRgate KRAB domain and setup of the proof of concept experiment

Supplementary Figure 2: Validation of single CD33 and CD13 targeting sgRNAs and CRISPRgate combination efficiency

Supplementary Figure 3: Off-target analysis of the 15 nt and 20 nt sgRNA targeting a fluorescent reporter.

Supplementary Figure 4: Validation of single BUB1 targeting sgRNAs and BUB1 depletion efficiency

Supplementary Figure 5: Staining and depletion validation for EMT and iPSCs

Supplementary Figure 6: CRISPRgate library design.

Supplementary Figure 7: Chimera rate and performance of the CRISPRgate library

Supplementary Figure 8: CRISPRgate achieves better reproducibility on read count and sgRNA LFC level

Supplementary Figure 9: CRISPRgate achieves better reproducibility on gene LFC level

Supplementary Figure 10: CRISPRgate demonstrates high sensitivity with increased consistency in sgRNA performance

Supplementary Figure 11: CRISPRgate achieves more significant hits on sgRNA and gene level

Supplementary Figure 12: CRISPRi efficiency depends on the TSS selection

Supplementary Tables S1-6.xlsx: Supplementary Table 1: Primers; Supplementary Table 2: CRISPRgate Library sgRNAs; Supplementary Table 3: Normalized screen count of all analyzed screens; Supplementary Table 4: sgRNA guide sequence; Supplementary Table 5: Normalized RNA-seq count and all DE-seq analysis; Supplementary Table 6: Amplicon sequencing count

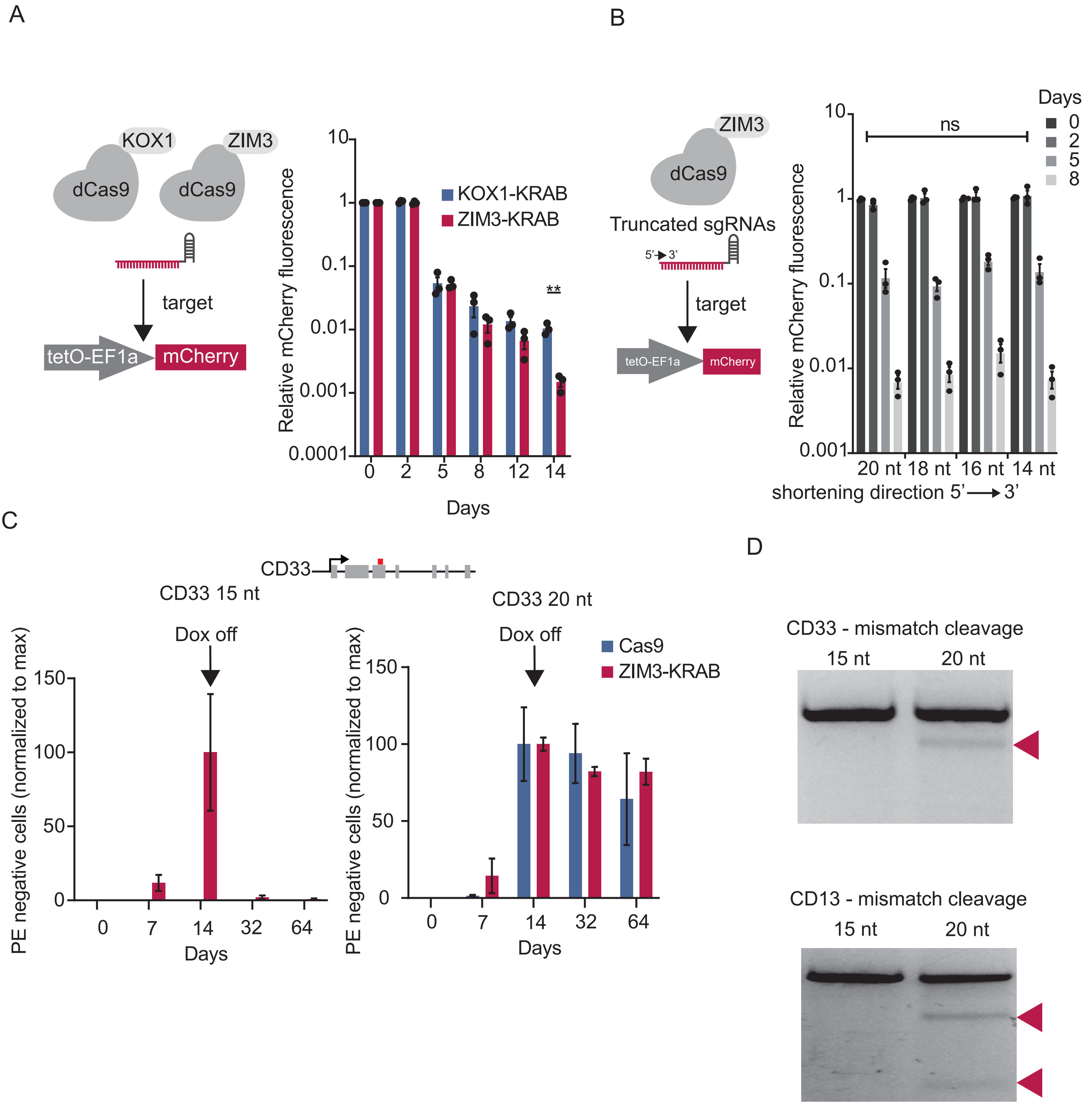
A
KOX1
ZIM3
dCas9
dCas9
target
tetO-EF1a
mCherry
Relative mCherry fluorescence
10
1
0.1
0.01
0.001
0.0001
KOX1-KRAB
ZIM3-KRAB
**
0
2
5
8
12
14
Days
B
ZIM3
dCas9
Truncated sgRNAs
5'
3'
target
tetO-EF1a
mCherry
Relative mCherry fluorescence
ns
Days
0
2
5
8
20 nt 18 nt 16 nt 14 nt
shortening direction 5' → 3'
C
CD33
CD33 15 nt
Dox off
PE negative cells (normalized to max)
150
100
50
0
0
7
14
32
64
Days
CD33 20 nt
Dox off
Cas9
ZIM3-KRAB
PE negative cells (normalized to max)
Days
D
CD33 - mismatch cleavage
15 nt
20 nt
CD13 - mismatch cleavage
15 nt
20 nt

**Supplementary Figure 1: Validation of the CRISPRgate KRAB domain and setup of the proof of concept experiment (A)** Flow-cytometric analysis of the mCherry expression in NIH/3T3 cells co-expressing the mCherry reporter, the dCas9-ZIM3 or dCas9-KOX1 fusion proteins and a sgRNA targeting the promoter region of the reporter. (n = 3, mean ± S.D..; **P ≤ 0.01, n.s. = non-significant; multiple t-tests with a two-stage set-up method of Benjamin to account for FDR). **(B)** Flow-cytometric analysis of the mCherry expression in NIH/3T3 cells co-expressing the mCherry reporter, the dCas9-ZIM3 fusion protein, and truncated sgRNAs (5' to 3') with the indicated length targeting the promoter region of the reporter. (n = 3, mean ± S.D..; n.s. = non-significant, ordinary two-way ANOVA with Tukey post-hoc test). **(C)** Experimental setup used to qualify sgRNA performance. CD13 and CD33 depletion was determined by immunostaining assay. **(D)** Quantification of CD33 expression by immunostaining of TF-1 erythroleukemia cells expressing Cas9 (blue) or ZIM3-Cas9 (red) and identical sgRNAs either 15 nt or 20 nt. ZIM3-Cas9/Cas9 expression was induced for 14 days and afterward. In case CD33 depletion was detected, on day 14 the percentage of CD33 negative cells was set as a reference, dox was removed and the recovery of the CD33 expression was monitored to determine the consistency of the induced genomic changes. (n = 3, mean ± S.D). **(E)** Representative gel image of a DNA mismatch cleavage assay at the end of the experiment determining non-homologous end-joining efficiency for the indicated sgRNAs at the CD13 and CD33 genes. Red arrowheads indicate cleavage products due to mismatches between WT CD13/CD33 DNA and Cas9-targeted CD13/CD33 genes.

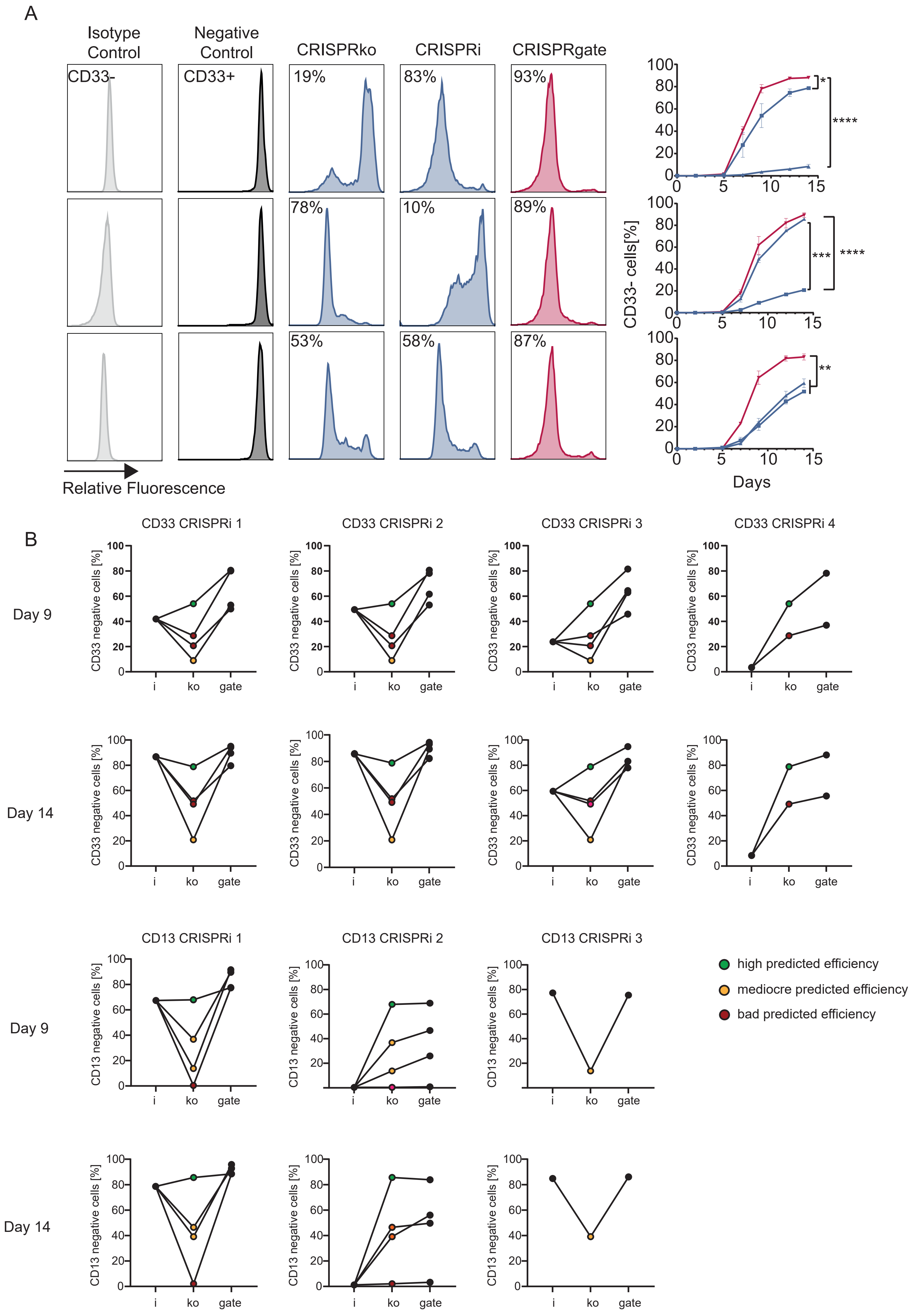
A
Isotype Control
Negative Control
CRISPRko
CRISPRi
CRISPRgate
CD33-
CD33+
19%
83%
93%
78%
10%
89%
53%
58%
87%
Relative Fluorescence
CD33- cells[%]
Days
*
****
***
****
**
B
CD33 CRISPRi 1
CD33 CRISPRi 2
CD33 CRISPRi 3
CD33 CRISPRi 4
Day 9
Day 14
CD33 negative cells [%]
i
ko
gate
CD13 CRISPRi 1
CD13 CRISPRi 2
CD13 CRISPRi 3
CD13 negative cells [%]
high predicted efficiency
mediocre predicted efficiency
bad predicted efficiency

**Supplementary Figure 2: Validation of single *CD33 and CD13* targeting sgRNAs and CRISPRgate combination efficiency. (A)** Flow cytometry quantification of the isotype and non-targeting control as well as for different *CD33* targeting sgRNAs and the respective CRISPRgate combination after 14 days as well as the overall time course of CD33 depletion (n = 3, mean ± s.e.m.; *P ≤ 0.05, **P ≤ 0.01, ***P ≤ 0.001, ****P ≤ 0.0001, n.s. = non-significant; two-way ANOVA with a Tukey post-hoc test). **(B)** CRISPR technology combination plot, median of the percentage of CD33/CD13 negative cells for the individual sgRNAs at day 9 and day 14 from figure 1E individually split for CRISPRi, CRISPRko, and the resulting indicated CRISPRgate combinations (n = 3).

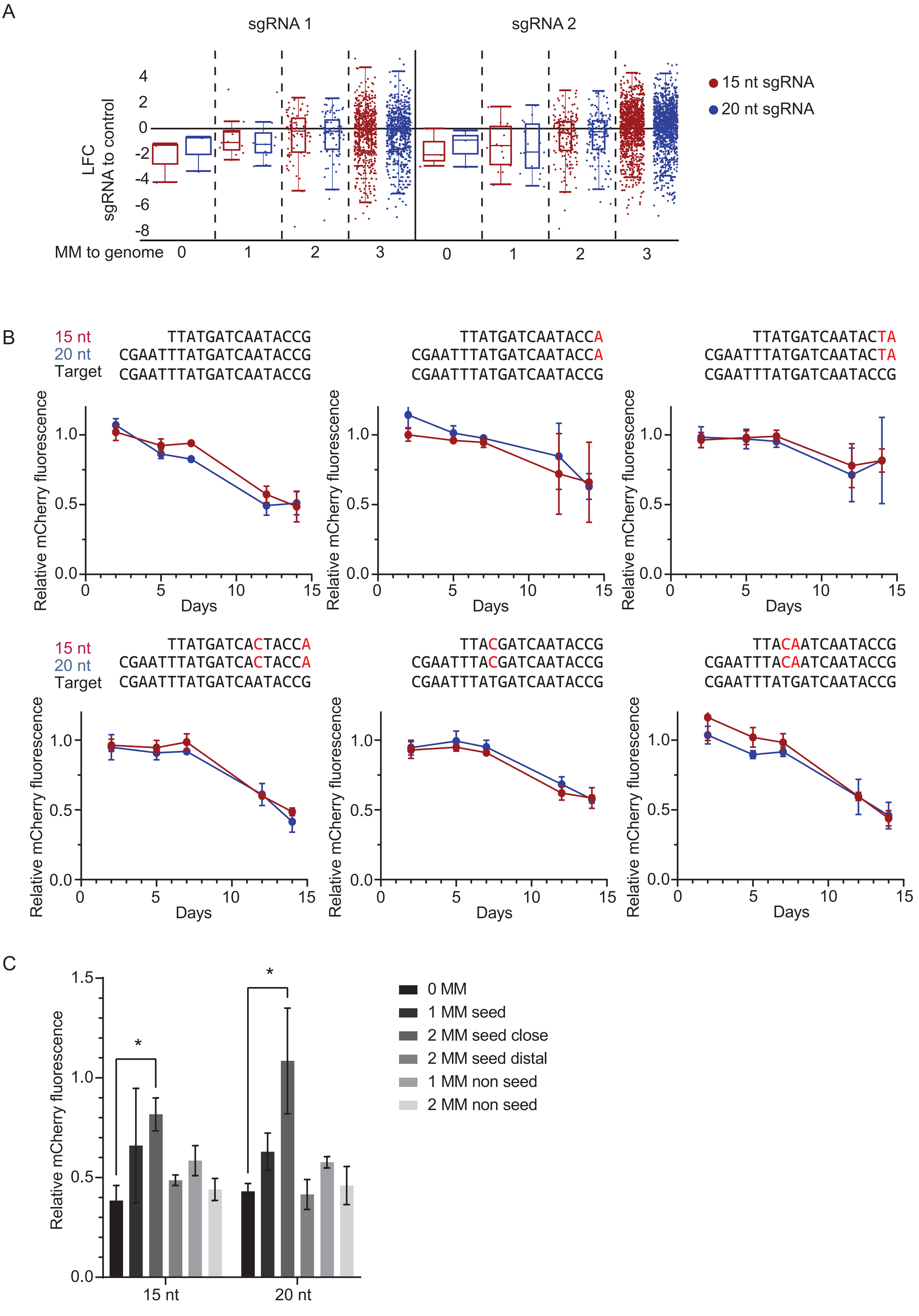
A
sgRNA 1
sgRNA 2
LFC
sgRNA to control
4
2
0
-2
-4
-6
-8
MM to genome
0
1
2
3
15 nt sgRNA
20 nt sgRNA
B
15 nt
20 nt
Target
TTATGATCAATACCG
CGAATTTATGATCAATACCG
CGAATTTATGATCAATACCG
TTATGATCAATACCA
CGAATTTATGATCAATACCA
CGAATTTATGATCAATACCG
TTATGATCAATACTA
CGAATTTATGATCAATACTA
CGAATTTATGATCAATACCG
TTATGATCACTACCA
CGAATTTATGATCACTACCA
CGAATTTATGATCAATACCG
TTACGATCAATACCG
CGAATTTACGATCAATACCG
CGAATTTATGATCAATACCG
TTACAATCAATACCG
CGAATTTACAATCAATACCG
CGAATTTATGATCAATACCG
Relative mCherry fluorescence
1.0
0.5
0.0
0
5
10
15
Days
C
*
1.5
1.0
0.5
0.0
Relative mCherry fluorescence
15 nt
20 nt
0 MM
1 MM seed
2 MM seed close
2 MM seed distal
1 MM non seed
2 MM non seed

**Supplementary figure 3: Off-target analysis of the 15 nt and 20 nt sgRNA targeting a fluorescent reporter. (A)** Log2 foldchange between the 15 nt or 20 nt CD33 targeting sgRNA from Figure 2 and the control sgRNA for all identified off-targets (up to 3 MM) of the 15 nt sgRNA split into groups by the MM number. **(B)** Flow-cytometric analysis of the mCherry expression in TF-1 cells co-expressing the mCherry reporter, the dCas9-ZIM3 as well as the indicated sgRNAs targeting the reporter. In red is indicated the respective off-target base towards the EF1a promoter expressing the mCherry. **(C)** Comparison of the relative mCherry fluorescence at day 14 between the sgRNA mismatches for the 15 nt sgRNA and the 20 nt sgRNA (n = 3, mean ± SEM.; *$P \leq 0.05$, **$P \leq 0.01$, ****$P \leq 0.0001$; two-way ANOVA with a Sidak post-hoc test).

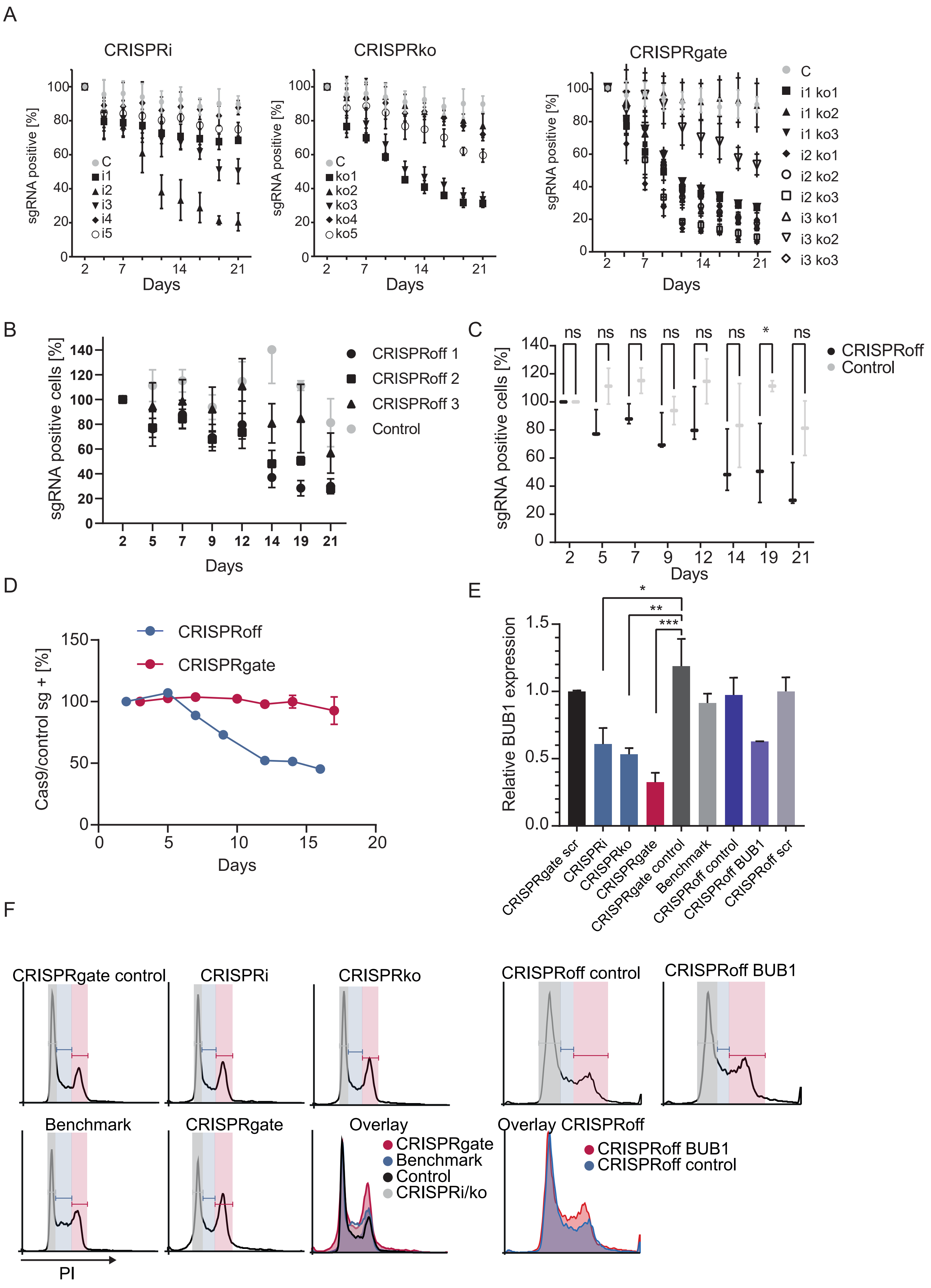

A
CRISPRi
CRISPRko
CRISPRgate
sgRNA positive [%]
Days
C
i1
i2
i3
i4
i5
ko1
ko2
ko3
ko4
ko5
i1 ko1
i1 ko2
i1 ko3
i2 ko1
i2 ko2
i2 ko3
i3 ko1
i3 ko2
i3 ko3
B
sgRNA positive cells [%]
CRISPRoff 1
CRISPRoff 2
CRISPRoff 3
Control
C
ns
*
CRISPRoff
Control
D
Cas9/control sg + [%]
CRISPRoff
CRISPRgate
E
Relative BUB1 expression
*
**
***
CRISPRgate scr
CRISPRi
CRISPRko
CRISPRgate
CRISPRgate control
Benchmark
CRISPRoff control
CRISPRoff BUB1
CRISPRoff scr
F
CRISPRgate control
CRISPRi
CRISPRko
CRISPRoff control
CRISPRoff BUB1
Benchmark
CRISPRgate
Overlay
CRISPRgate
Benchmark
Control
CRISPRi/ko
Overlay CRISPRoff
CRISPRoff BUB1
CRISPRoff control
PI

**Supplementary figure 4: Validation of single *BUB1* targeting sgRNAs and BUB1 depletion efficiency. (A)** Raw data from figure 3A showing the individual competitive proliferation assays of TF-1 cells expressing the indicated single sgRNAs as well as the respective CRISPRgate combinations targeting *BUB1*. Shown is the relative fraction of GFP+/sgRNA+ cells relative to the initial measurement over the course of 21 days. (n = 3, mean ± s.e.m.) **(B)** Raw data of the individual dual CRISPRoff sgRNAs targeting *BUB1* and the control targeting a non-essential gene. **(C)** Summary of the proliferative effect of *BUB1* targeting sgRNAs for CRISPRoff and the respective control targeting a non-essential gene. **(D)** Percentage of cells co-expressing CRISPRoff/CRISPRgate and a non-targeting sgRNA control. **(E)** Relative *BUB1* expression of HEK293 cells expressing sgRNAs targeting *BUB1* normalized to b-actin and a neutral control sgRNA to validate the improved CRISPRgate effect observed in the proliferation assays. (n = 3, mean ± S.D..; *P ≤ 0.05, **P ≤ 0.01, ***P ≤ 0.001, n.s. = non-significant; two-way ANOVA with a Dunnet post-hoc test). **(F)** Exemplary histograms of the P.I. staining of the individual samples and the corresponding distribution in G1- (grey), S- (blue), and G2/M-phase (red). An overlay of all histograms was plotted to further highlight the difference in cell cycle distribution.

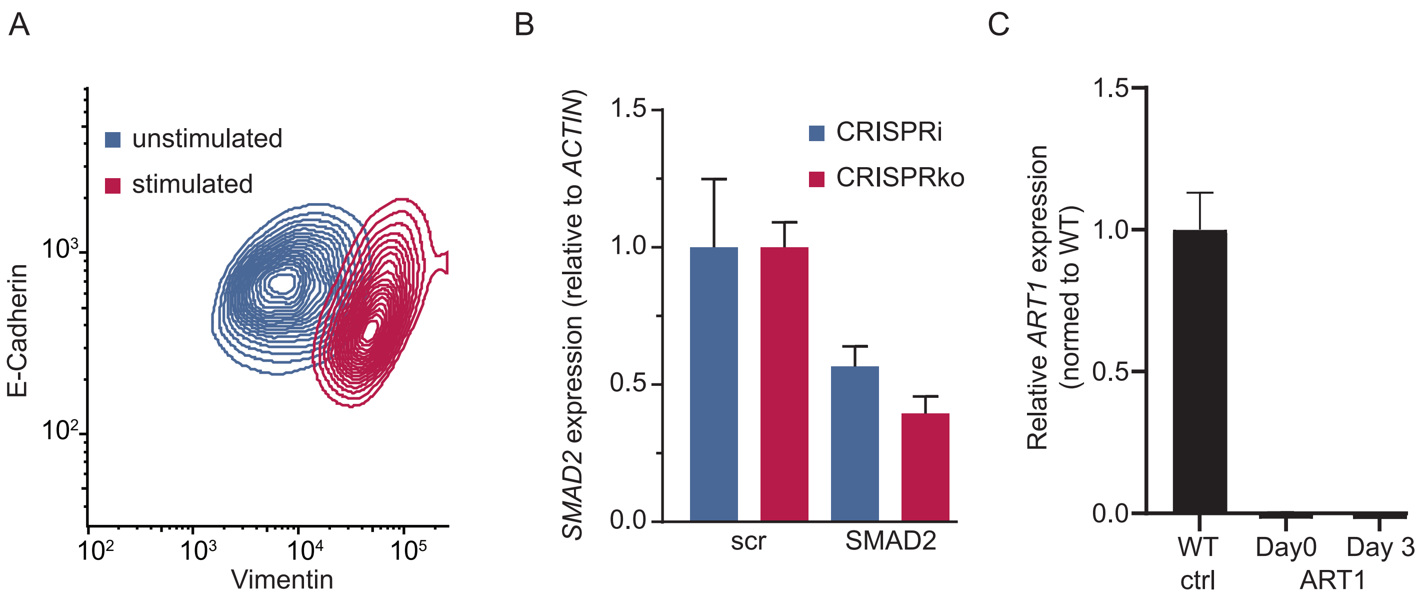
A
unstimulated
stimulated
E-Cadherin
Vimentin
B
CRISPRi
CRISPRko
SMAD2 expression (relative to ACTIN)
scr
SMAD2
C
Relative ART1 expression (normed to WT)
WT ctrl
Day0
Day 3
ART1

**Supplementary Figure 5: Staining and depletion validation for EMT and iPSCs. (A)** MCF10A cells were stained with antibodies binding to the intracellular proteins Vimentin and E-Cadherin used to identify and gate for the epithelial cell population (blue) and mesenchymal cell population (red). **(B)** Relative *SMAD2* expression of MCF10A cells expressing sgRNAs targeting a non-essential control or *SMAD2* normalized to b-actin to validate the loss of SMAD2. **(C)** Relative ART1 expression of unstimulated and stimulated iPSC cells expressing CRISPRgate and a sgRNA combination targeting ART1 normalized to *b-Actin* and WT.

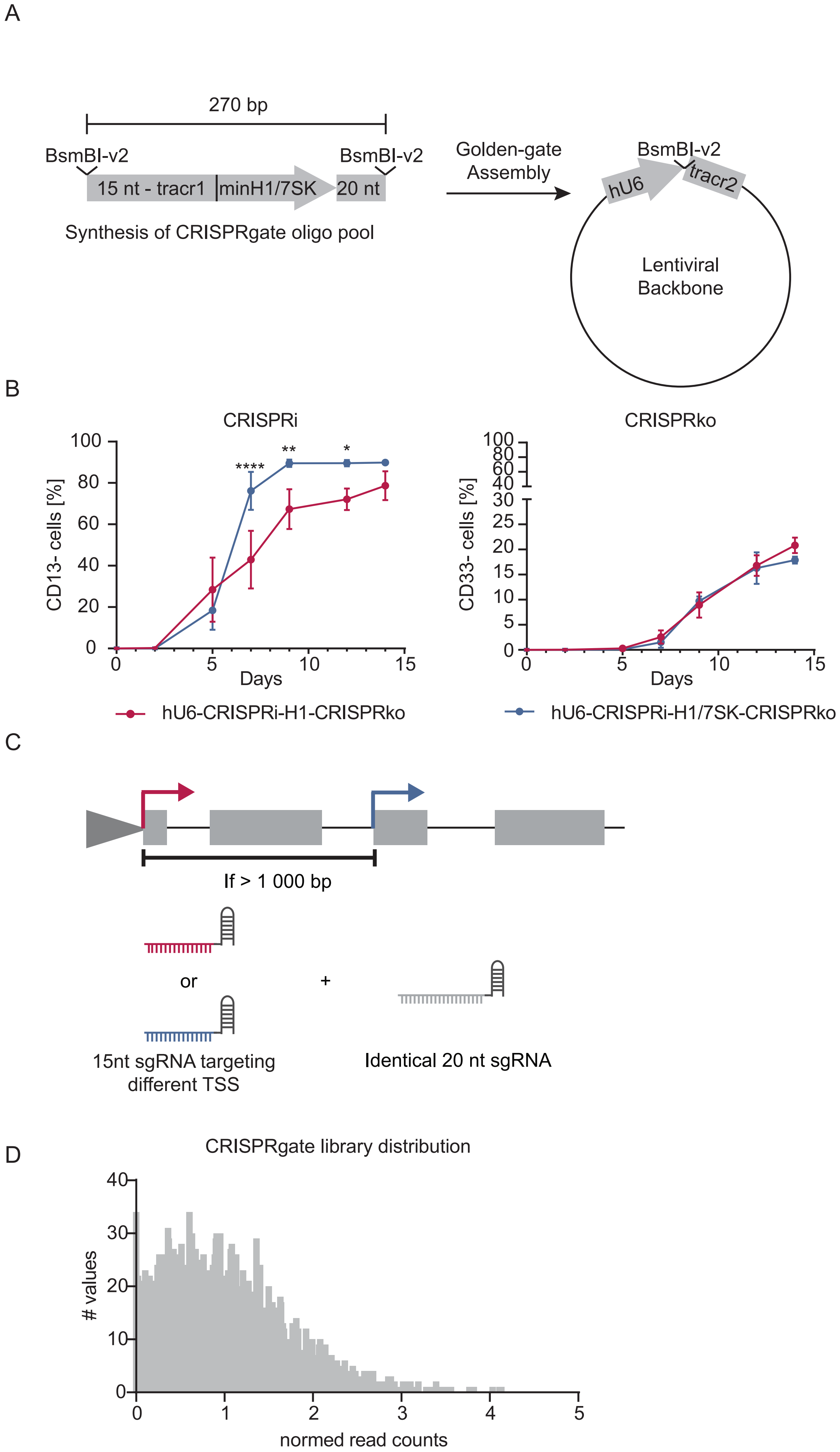
A
270 bp
BsmBI-v2
BsmBI-v2
15 nt - tracr1
minH1/7SK
20 nt
Synthesis of CRISPRgate oligo pool
Golden-gate Assembly
BsmBI-v2
hU6
tracr2
Lentiviral Backbone
B
CRISPRi
CRISPRko
CD13- cells [%]
CD33- cells [%]
Days
Days
****
**
*
hU6-CRISPRi-H1-CRISPRko
hU6-CRISPRi-H1/7SK-CRISPRko
C
If > 1 000 bp
or
+
15nt sgRNA targeting different TSS
Identical 20 nt sgRNA
D
CRISPRgate library distribution
### values
normed read counts

**Supplementary Figure 6: CRISPRgate library design.** **(A)** Oligo design of the CRISPRgate library and the subsequent cloning strategy into the lentiviral backbone. **(B)** Time-resolved quantification of dual CD13 and CD33 depletion in TF-1 cells using the hU6 promoter to express identical CD13 sgRNAs as well as the H1 or minimal H1/7SK promoter to express the CD33 targeting sgRNA (n = 3, mean ± S.D.; *P ≤ 0.05, **P ≤ 0.01, ****P ≤ 0.0001; two-way ANOVA with a Sidak post-hoc test). **(C)** CRISPRgate combination strategy for genes harbouring more than one TSS with a distance of > 1000 bp. **(D)** CRISPRgate sgRNA library distribution, measured and calculated by deep-sequencing.

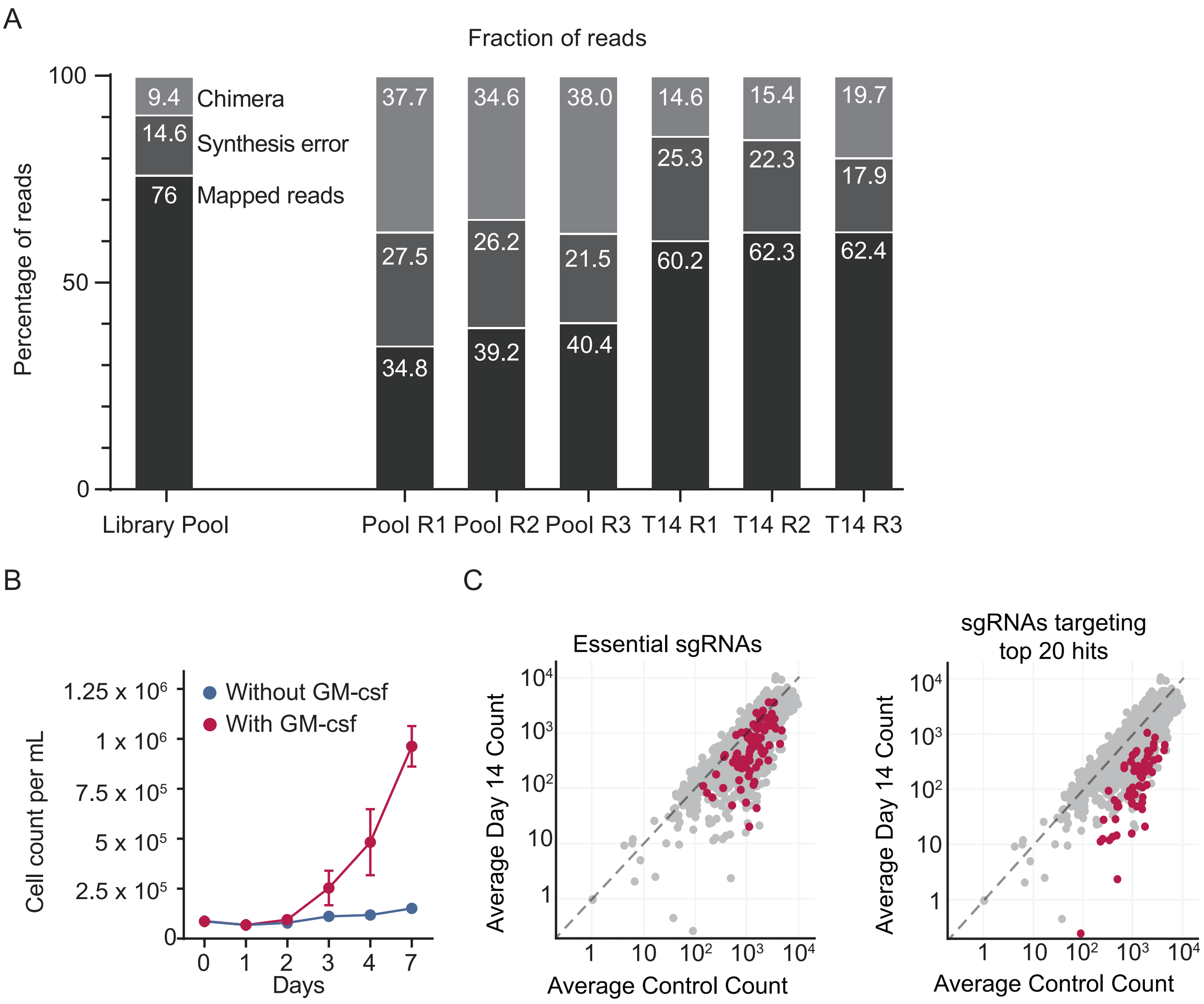
A
Fraction of reads
Percentage of reads
100
50
0
9.4
Chimera
14.6
Synthesis error
76
Mapped reads
37.7
27.5
34.8
34.6
26.2
39.2
38.0
21.5
40.4
14.6
25.3
60.2
15.4
22.3
62.3
19.7
17.9
62.4
Library Pool
Pool R1
Pool R2
Pool R3
T14 R1
T14 R2
T14 R3
B
Cell count per mL
Without GM-csf
With GM-csf
Days
C
Essential sgRNAs
sgRNAs targeting top 20 hits
Average Day 14 Count
Average Control Count

**Supplementary Figure 7: Chimera rate and reproducibility of the CRISPRgate library. (A)** Attributed mapped reads, synthesis error reads, and the chimera rate of the oligo pool replicates and the day 14 replicates of the CRISPRgate screen after library preparation and sequencing. **(B)** Proliferation of TF-1 cells grown in RPMI media supplemented with or without the growth factor GM-CSF. **(C)** Left: normalized read counts of the sgRNAs targeting essential genes for the library pool after 14 days of screening. Right: normalized read counts of sgRNAs targeting the top 20 gene hits identified in the CRISPRgate screen of the library pool compared to day 14 after induction of ZIM3-Cas9 expression.

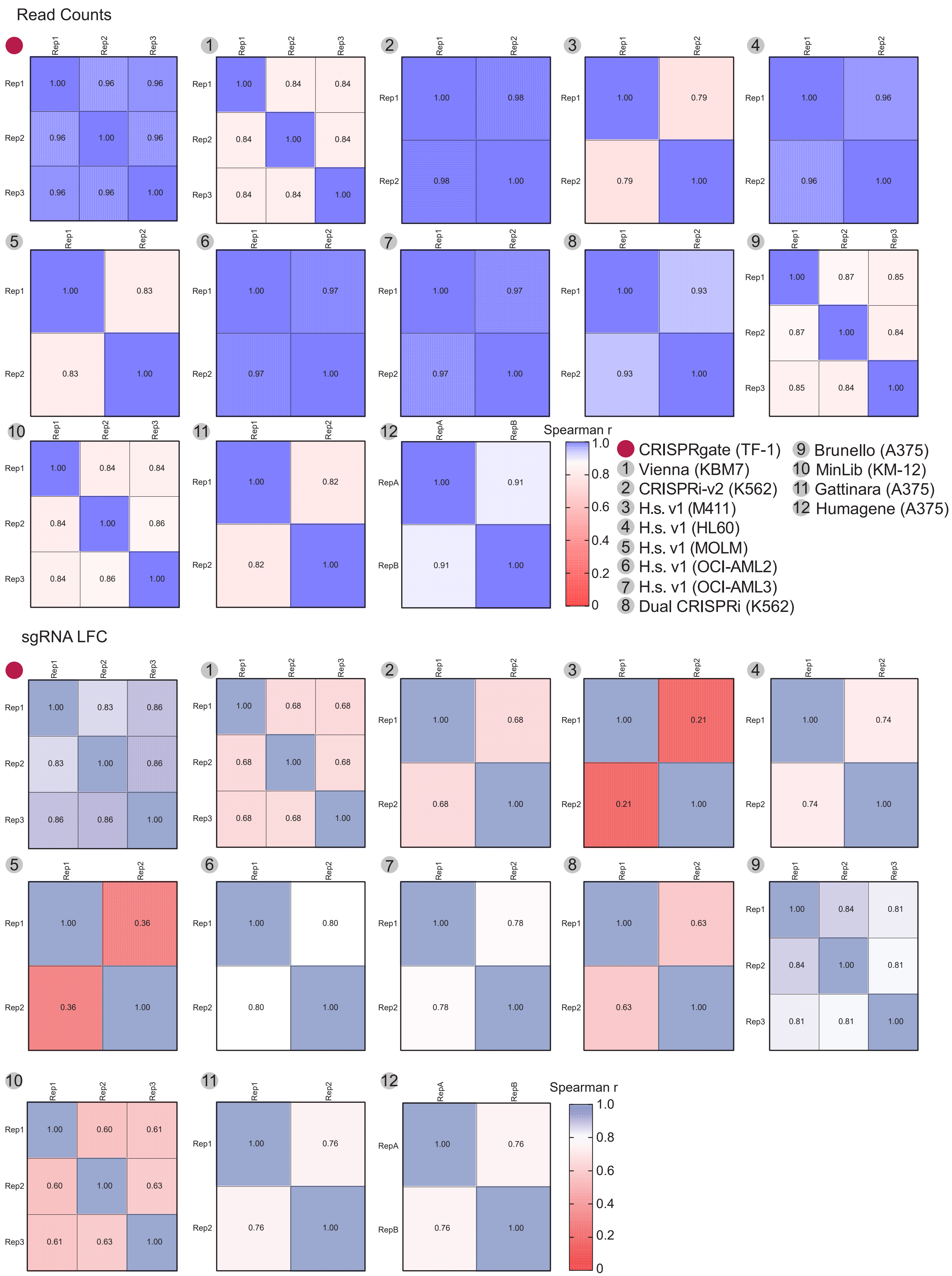

Read Counts
sgRNA LFC
Spearman r
CRISPRgate (TF-1)
1 Vienna (KBM7)
2 CRISPRi-v2 (K562)
3 H.s. v1 (M411)
4 H.s. v1 (HL60)
5 H.s. v1 (MOLM)
6 H.s. v1 (OCI-AML2)
7 H.s. v1 (OCI-AML3)
8 Dual CRISPRi (K562)
9 Brunello (A375)
10 MinLib (KM-12)
11 Gattinara (A375)
12 Humagene (A375)

**Supplementary Figure 8: CRISPRgate achieves better reproducibility on read count and sgRNA LFC level.** Heat maps for each CRISPR screen analyzed in Figure 4D. Top: Spearman correlation of read counts of individual screening replicates for each analyzed screen. Bottom: Spearman correlation of the sgRNA LFC of individual screening replicates for each analyzed screen.

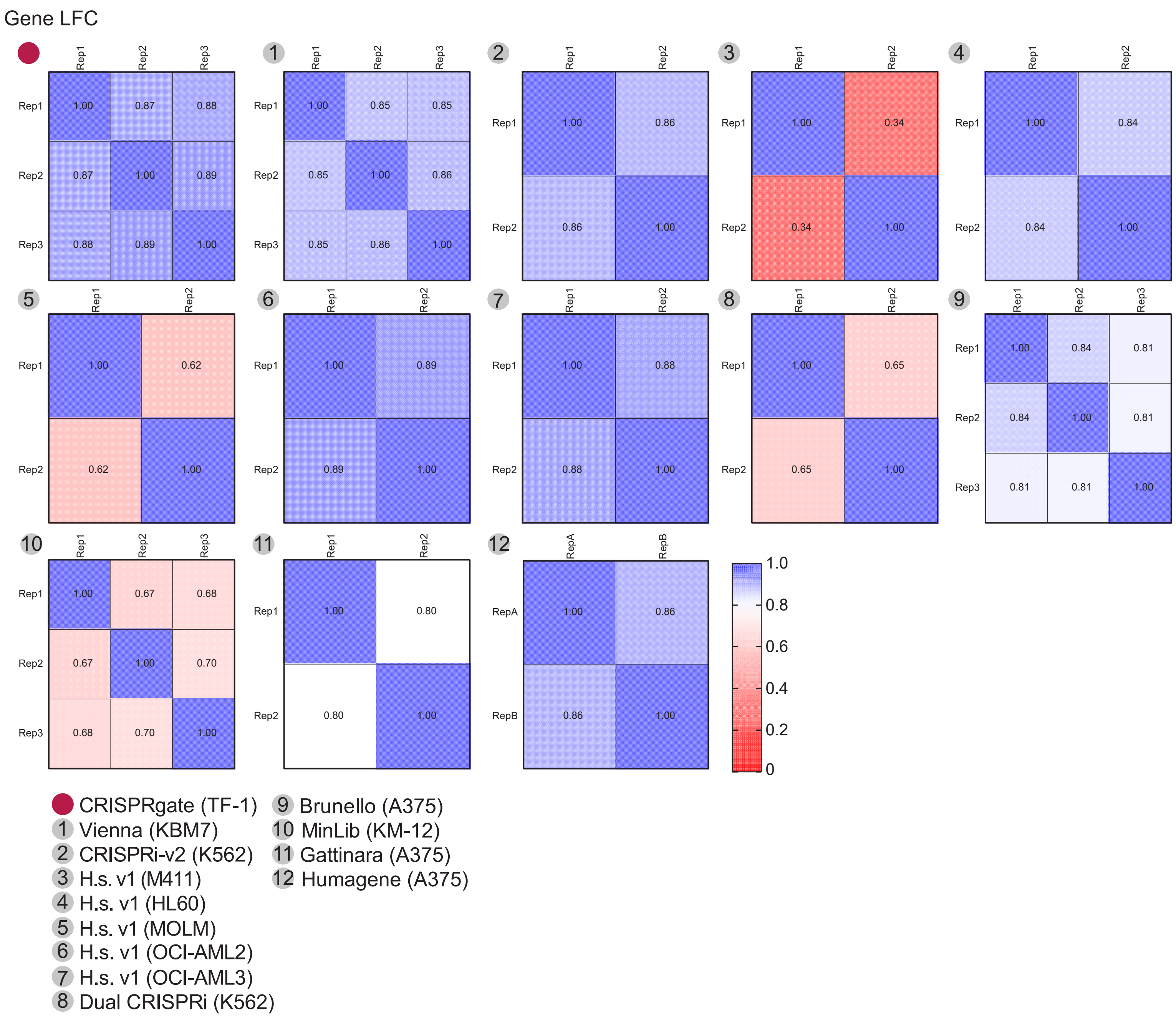
Gene LFC
1.00 0.87 0.88
0.87 1.00 0.89
0.88 0.89 1.00
1.00 0.85 0.85
0.85 1.00 0.86
0.85 0.86 1.00
1.00 0.86
0.86 1.00
1.00 0.34
0.34 1.00
1.00 0.84
0.84 1.00
1.00 0.62
0.62 1.00
1.00 0.89
0.89 1.00
1.00 0.88
0.88 1.00
1.00 0.65
0.65 1.00
1.00 0.84 0.81
0.84 1.00 0.81
0.81 0.81 1.00
1.00 0.67 0.68
0.67 1.00 0.70
0.68 0.70 1.00
1.00 0.80
0.80 1.00
1.00 0.86
0.86 1.00
Rep1
Rep2
Rep3
RepA
RepB
1.0
0.8
0.6
0.4
0.2
0
CRISPRgate (TF-1)
1 Vienna (KBM7)
2 CRISPRi-v2 (K562)
3 H.s. v1 (M411)
4 H.s. v1 (HL60)
5 H.s. v1 (MOLM)
6 H.s. v1 (OCI-AML2)
7 H.s. v1 (OCI-AML3)
8 Dual CRISPRi (K562)
9 Brunello (A375)
10 MinLib (KM-12)
11 Gattinara (A375)
12 Humagene (A375)

**Supplementary Figure 9: CRISPRgate achieves better reproducibility on gene LFC level.** Heat maps for each CRISPR screen analysed in Figure 4D in which the gene LFC of individual screening replicates per screen were correlated using the spearman correlation analysis.

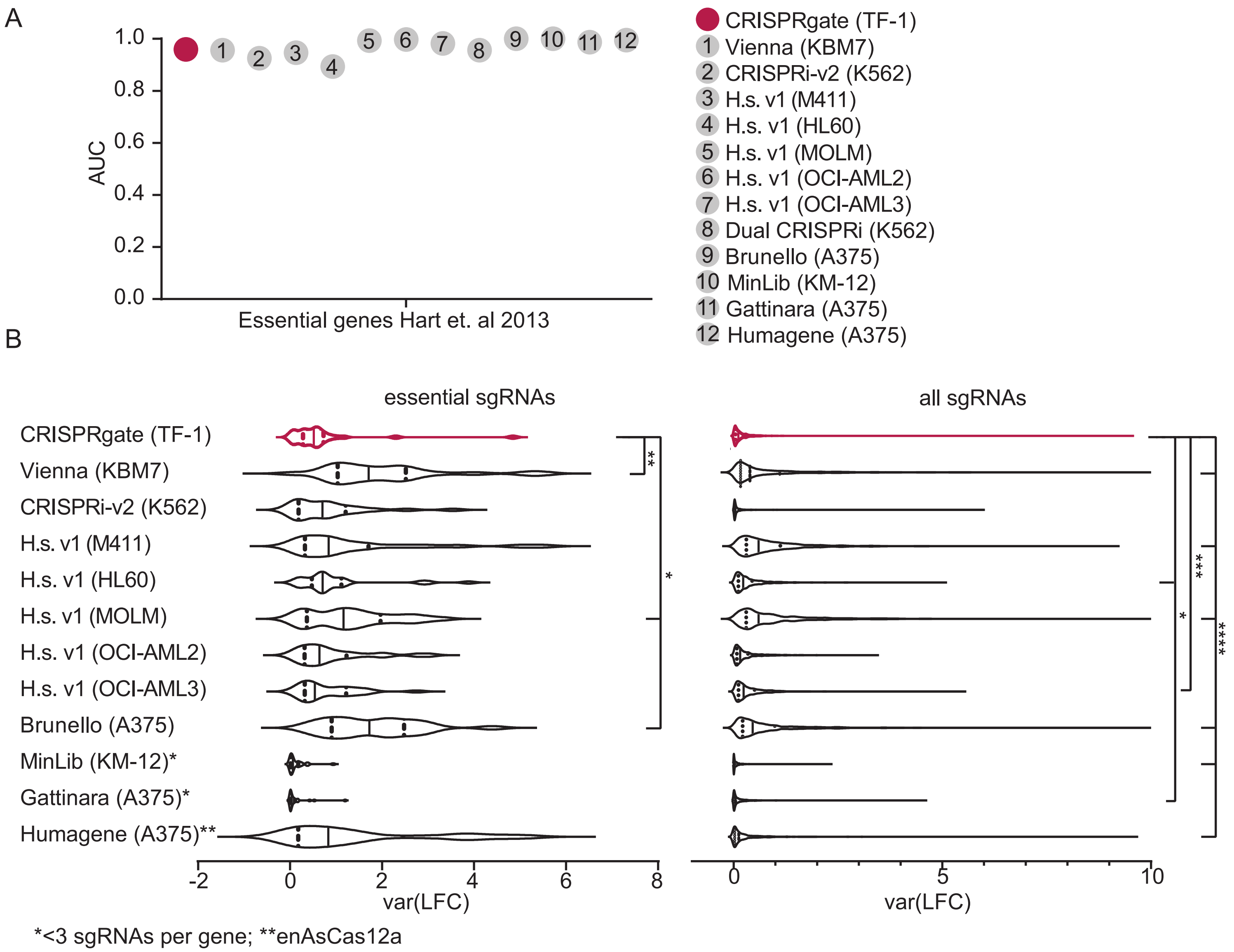
A
1.0
0.8
0.6
0.4
0.2
0.0
AUC
Essential genes Hart et. al 2013
CRISPRgate (TF-1)
1 Vienna (KBM7)
2 CRISPRi-v2 (K562)
3 H.s. v1 (M411)
4 H.s. v1 (HL60)
5 H.s. v1 (MOLM)
6 H.s. v1 (OCI-AML2)
7 H.s. v1 (OCI-AML3)
8 Dual CRISPRi (K562)
9 Brunello (A375)
10 MinLib (KM-12)
11 Gattinara (A375)
12 Humagene (A375)
B
essential sgRNAs
all sgRNAs
CRISPRgate (TF-1)
Vienna (KBM7)
CRISPRi-v2 (K562)
H.s. v1 (M411)
H.s. v1 (HL60)
H.s. v1 (MOLM)
H.s. v1 (OCI-AML2)
H.s. v1 (OCI-AML3)
Brunello (A375)
MinLib (KM-12)*
Gattinara (A375)*
Humagene (A375)**
**
*
***
*
****
-2
0
2
4
6
8
var(LFC)
0
5
10
var(LFC)
*<3 sgRNAs per gene; **enAsCas12a

**Supplementary Figure 10: CRISPRgate demonstrates high sensitivity with increased consistency in sgRNA performance. (A)** The area under the curve **(**AUC) of ROC curves based on essential and non-essential genes for the CRISPRgate screen and other published screens shown in Figure 4C. **(B)** Violin plots comparing the variance among sgRNAs targeting the same gene observed with the CRISPRgate system with a set of published CRISPR screening approaches. Left: Comparison of the variance of sgRNAs targeting the same genes from a subset of essential genes. Right: Comparison of the variance of sgRNAs targeting the same genes for all genes investigated in the CRISPRgate screen.

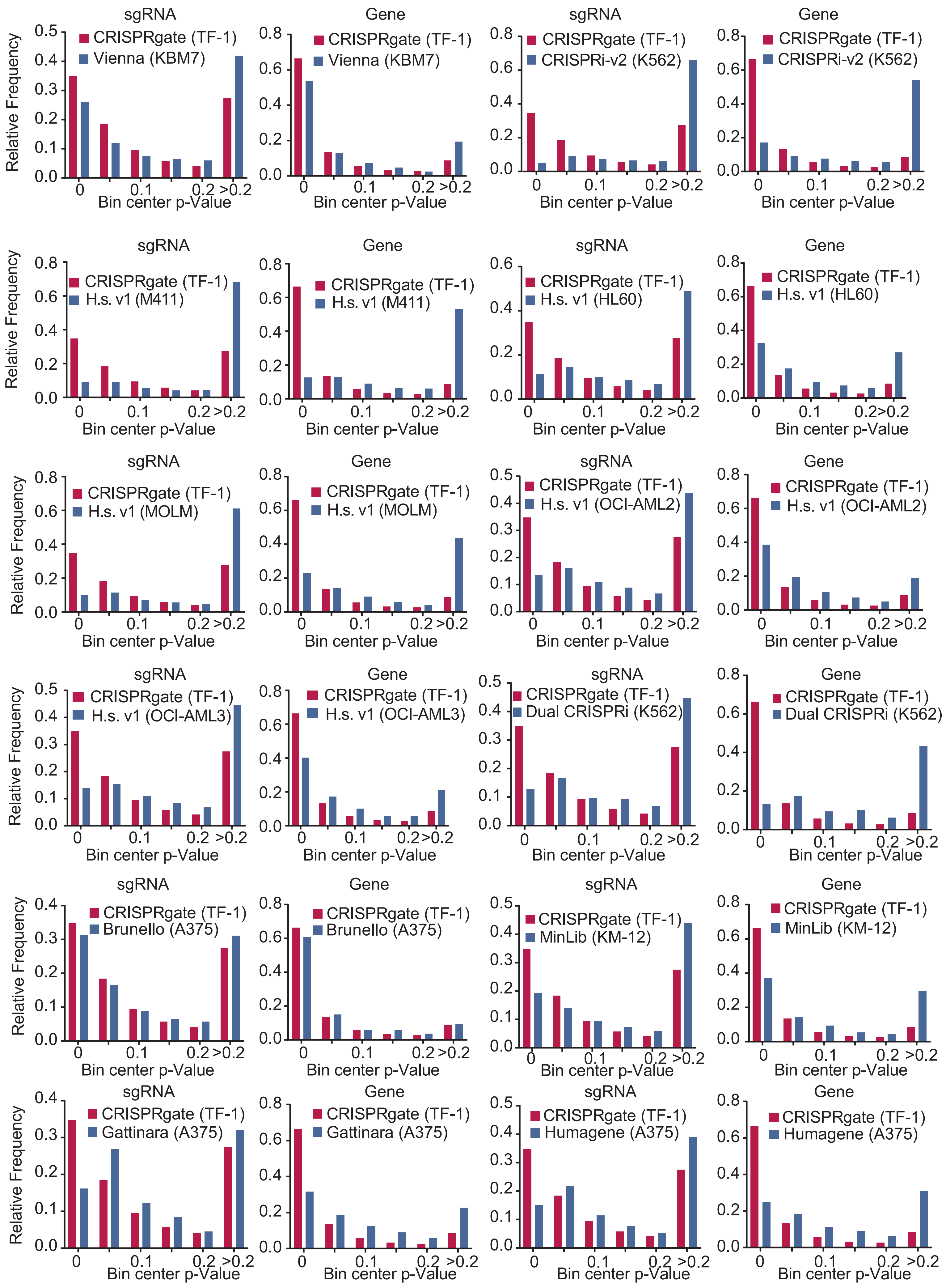

sgRNA
Gene
Relative Frequency
Bin center p-Value
CRISPRgate (TF-1)
Vienna (KBM7)
CRISPRi-v2 (K562)
H.s. v1 (M411)
H.s. v1 (HL60)
H.s. v1 (MOLM)
H.s. v1 (OCI-AML2)
H.s. v1 (OCI-AML3)
Dual CRISPRi (K562)
Brunello (A375)
MinLib (KM-12)
Gattinara (A375)
Humagene (A375)

**Supplementary Figure 11: CRISPRgate achieves more significant hits on sgRNA and gene level.** Histograms comparing the relative p-value frequencies of sgRNAs as well as the p-value for each gene in the CRISPRgate screen with a set of published CRISPR screening approaches.

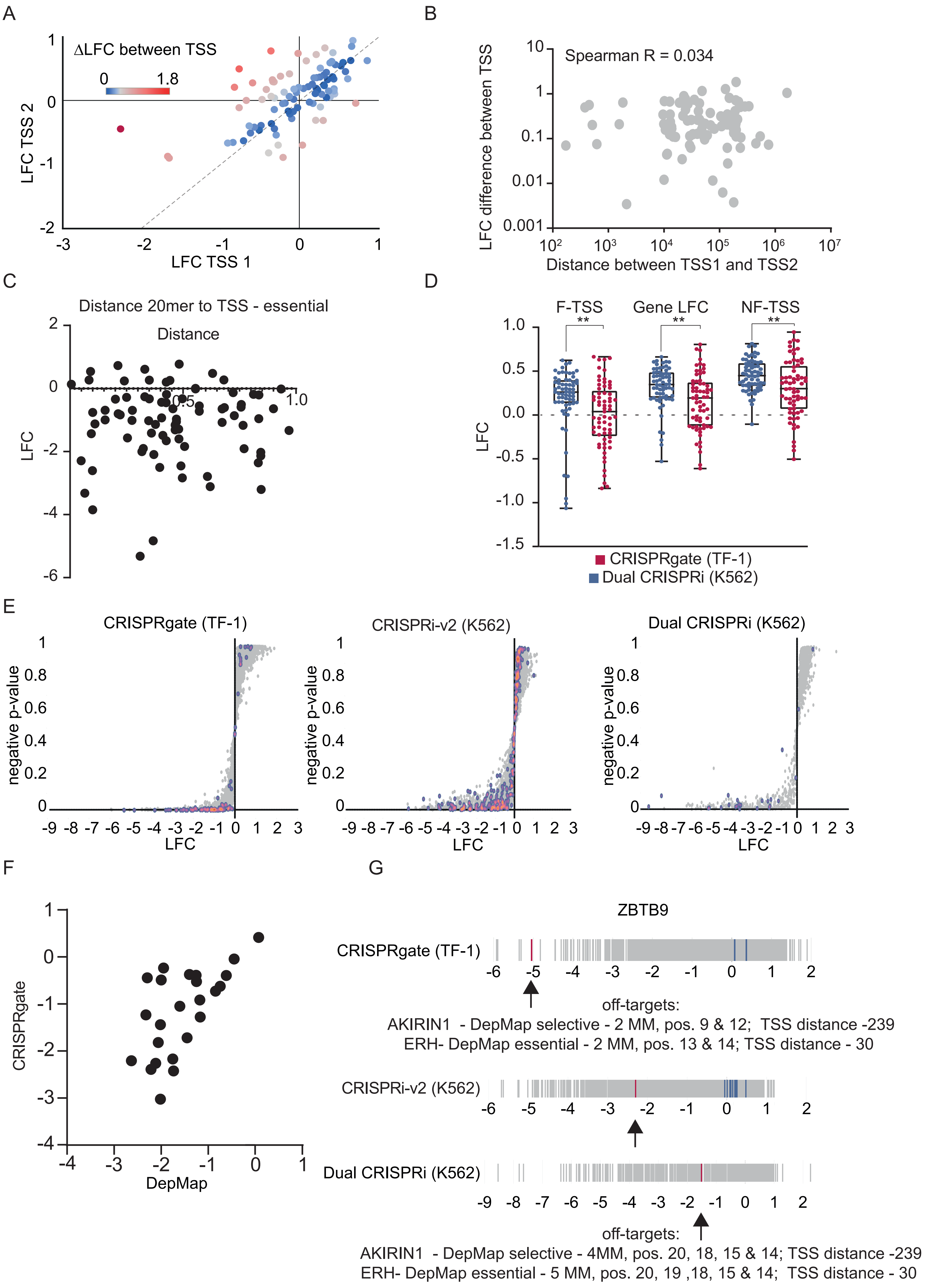

A
ΔLFC between TSS
0
1.8
LFC TSS 2
LFC TSS 1
B
Spearman R = 0.034
LFC difference between TSS
Distance between TSS1 and TSS2
C
Distance 20mer to TSS - essential
Distance
LFC
D
F-TSS
Gene LFC
NF-TSS
LFC
CRISPRgate (TF-1)
Dual CRISPRi (K562)
E
CRISPRgate (TF-1)
CRISPRi-v2 (K562)
Dual CRISPRi (K562)
negative p-value
LFC
F
CRISPRgate
DepMap
G
ZBTB9
CRISPRgate (TF-1)
off-targets:
AKIRIN1 - DepMap selective - 2 MM, pos. 9 & 12; TSS distance -239
ERH- DepMap essential - 2 MM, pos. 13 & 14; TSS distance - 30
CRISPRi-v2 (K562)
Dual CRISPRi (K562)
off-targets:
AKIRIN1 - DepMap selective - 4MM, pos. 20, 18, 15 & 14; TSS distance -239
ERH- DepMap essential - 5 MM, pos. 20, 19 ,18, 15 & 14; TSS distance - 30

**Supplementary Figure 12: CRISPRi efficiency depends on the TSS selection. (A)** Scatter plot depicting the LFC of genes which have more than one TSS and are 1 000 bp distant from each other split into the cumulative LFC for sgRNAs targeting one TSS and sgRNAs targeting the other. The ΔLFC was calculated between the two TSSs and is displayed with a blue-to-red color gradient depending on the ΔLFC. **(B)** Relationship between the ΔLFC of two different TSS of the same gene and the distance between both TSSs. **(C)** Distance of the CRISPRko sgRNA to the TSS plotted against the LFC to identify if the CRISPRko sgRNA is also affecting gene expression through the ZIM3 domain during Cas9 binding and cleavage. **(D)** The difference in LFC between the CRISPRgate screen and the dual CRISPRi screen when targeting only the functional (F) TSS, the non-functional (NF) TSS, or the cumulative LFC for each gene. **(E)** Plot of the p-value for each sgRNA over the LFC of the sgRNA for CRISPRgate, a CRISPRi screen (16) and a dual CRISPRi screen (28). The sgRNAs targeting essential genes are displayed as density plot in red. **(F)** Plot of the CRISPRgate LFC over the DepMap score for all essential genes targeted in the CRISPRgate screen. **(G)** LFC for the CRISPRgate sgRNA combinations targeting *ZBTB9*.
